# Unlocking the genomic repertoire of a cultivated megaphage

**DOI:** 10.1101/2024.12.16.628780

**Authors:** Andra Buchan, Stephanie Wiedman, Kevin Lambirth, Madeline Bellanger-Perry, Jose L. Figueroa, Elena T. Wright, Patil Shivprasad Suresh, Qibin Zhang, Julie A. Thomas, Philip Serwer, Richard Allen White

## Abstract

Megaphages are bacteriophages (i.e., phages) with exceptionally large genomes that are ecosystem cosmopolitans, infect various bacterial hosts, and have been discovered across various metagenomic datasets globally. To date, almost all megaphages have evaded cultivation, with only phage G being in active culture for over 50 years. We examined with multiomics this five decades long cultivated history from nine different laboratories with five different lab variants to the modern era. In this work, we resolved the five complete phage G genomes, the particle proteome, *de novo* methylome, and used artificial intelligence (AI) to annotate the genome of phage G. Phage G is one of the largest phages with a size of >0.6 µm, about half the width of the host cell, and a 499 kbp, non-permuted, linear genome that has, uniquely among known phages, two pairs of ends. Its closest known relative is Moose phage W30-1 which was metagenomically assembled without cultivation from a moose rumen sample. Phage G has >650 protein-coding open reading frames (ORFs), with >65% being hypothetical proteins with no known function, with the rest of the genome geared towards nucleic acid replication (e.g., helicases, polymerases, endonucleases) and are structural in nature (e.g., capsid, tail, portal, terminase). The genome encodes a 35 kbp stretch with 66 ORFs without any known functional homology, a cryptic genomic region that is roughly the size of phage lambda. Phage G has an expansive repertoire of auxiliary metabolic genes (AMGs) acquired from its bacterial host, including a *phoH*,*ftsZ*,*UvsX/RecA-like, gyrA, gyrB*,and *DHFR*. Furthermore, AMGs discovered in phage G could manipulate host sporulation (*sspD, RsfA, spoK*) and antiviral escape genes (e.g., anti-CBass nuclease and Anti-Pycsar protein). Phage proteomics found >15% of the protein ORFs were present in either the wild-type or mutant variants of phage G, including genes involved in replication (e.g.,*UvsX/RecA-like*), host sporulation, as well as structural genes (e.g., capsid, tail, portal). The methylome of phage G was localized to the cryptic region with limited functional homology, with supervised machine learning (i.e., HMMs) was unable to resolve this region, but was resolved with protein structural AI. This cryptic region was a hot spot for methylation at 32%, where many of the functions of the ORF are still unknown. Our study represents a doorway into the complexity of the genomic repertoire of the only cultivated megaphage, highlighting five decades of continuous cultivation for the first time.

## Introduction

Phage are ubiquitous, cosmopolitan, and present within all of Earth’s diverse ecosystems (e.g., hot springs, soils, animal guts, wastewater, microbialites, and marine ecosystems), but the majority have genomes that are less than 200 kbp (Al-Shayeb et al., 2020; Breitbart et al., 2004; Carreira et al., 2024; Cook et al., 2024; Devoto et al., 2019; Michniewski et al., 2021; White III et al., 2020; 2021; Weinheimer et al., 2022). Megaphages (i.e., phages with genomes 500 kbp or larger) have been metagenomic detected across diverse ecosystems from animal guts, wastewater, and oceans (Al-Shayeb et al., 2020; Cook et al., 2024; Devoto et al., 2019; Michniewski et al., 2021; Weinheimer et al., 2022). Only one megaphage has been physically isolated and cultivated, that is phage G in 1968 (Ageno et al., 1973; Donelli, 1968; Donelli et al., 1975; 1976) (**Fig 1**). A draft genome was released to GenBank by Roger Hendrix *et al*.at the University of Pittsburgh in 2012 (GenBank ID: NC_023719.1,**Fig 1**). Commonly, megaphages found in the guts of animals, including humans, are known as Lak Phages (Cook et al., 2024). The largest phage genome discovered from metagenomics was 735 kbp, isolated from Lac Pavin, a freshwater meromictic crater lake in France (Al-Shayeb et al., 2020). Another megaphage recently found via metagenomics was Mar_Mega_1 from marine waters within Plymouth Sound, UK (Michniewski et al., 2021). However, in 50 years, no megaphage has ever been cultivated beyond phage G.

**Figure 1:**
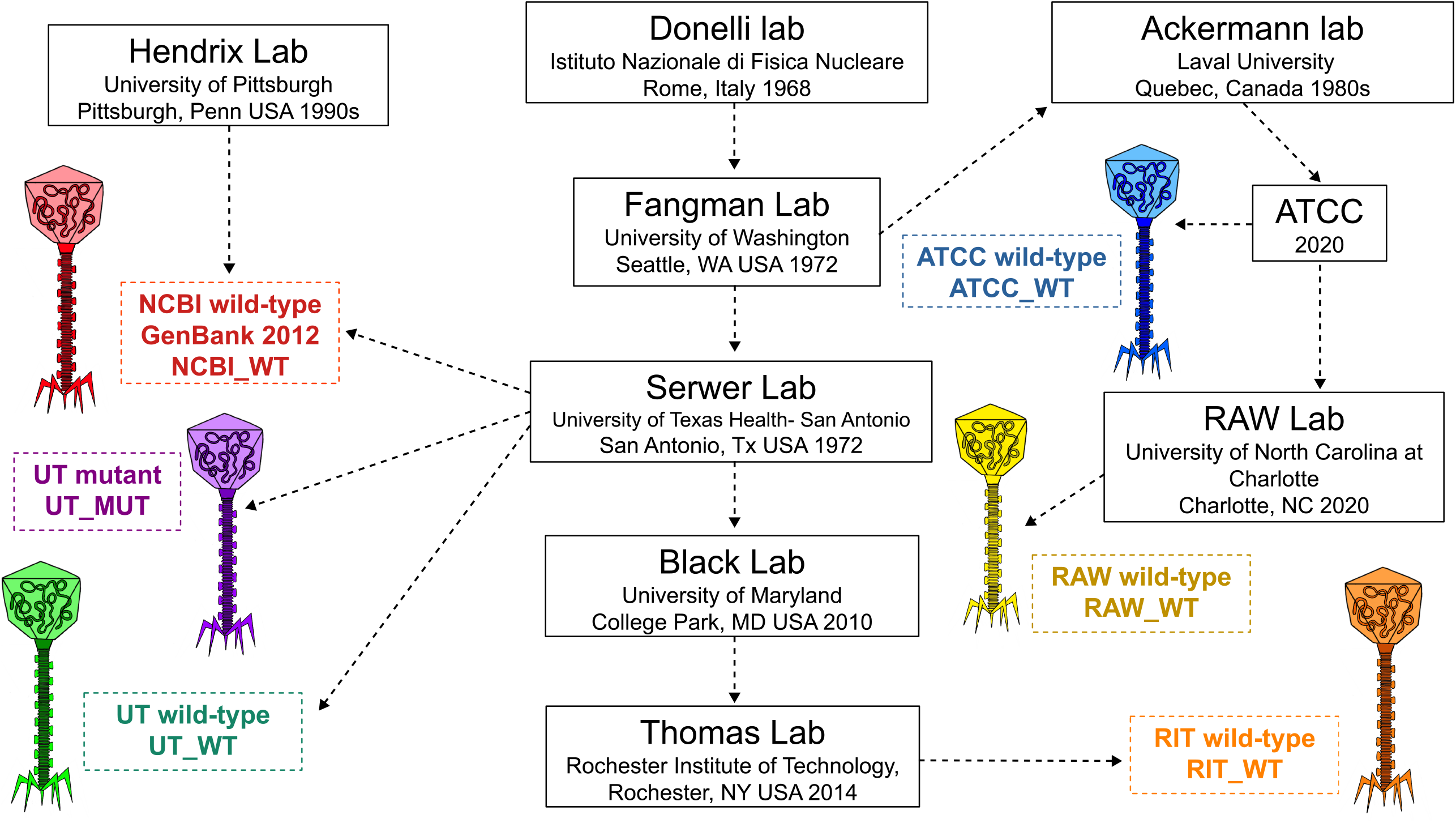
The cultivation history of phage G. This provides the cultivation history from its isolation in the Donelli lab (1968), through eight other laboratories, to the modern day over a span of five decades.

The history of phage G starts over 50 years ago at the Donelli lab at *Istituto di Fisica* in Rome (Donelli, 1968; Donelli et al., 1975; 1976) (**Fig 1)**.The host for phage G was recently revised to *Lysinibacillus* (González et al., 2020). The original environmental origin of phage G is unknown. Phage G was transferred to the lab of W.F. Fangman at the University of Washington, Seattle, WA in the 1970s, which sent phage G to the laboratory of Philip Serwer (**Fig 1**).

Unknown until 2020, the H.W. Ackermann-laboratory, a lab that personally imaged over 5,500 phages, used phage G as a reference for TEM and then donated it posthumously to the American Type Culture Collection (ATCC) in 2020. The Ackermann-laboratory had also received phage G from the Fangman-laboratory, likely in the 1980s or 1990s (Ackermann, 2007;**Fig 1**).

As mentioned above, the Serwer-laboratory sent R.W. Hendrix (RWH) the wild-type strain (which we called here the UT_WT, University of Texas - Wild Type) for genome sequencing in approximately 2001. We call this strain the NCBI wild-type, derived from UT_WT (NCBI_WT,**Fig 1**). RWH noticed that, at some point in its propagation, NCBI_WT was able to grow well in liquid media, unlike the original UT_WT strain, suggesting that a spontaneous mutation had occurred. The Serwer-laboratory deliberately isolated a clearer plaque mutant (i.e., UT_MUT) (**Fig 1**). The Serwer-laboratory sent the original UT_WT to Lindsey Black and Julie A. Thomas in 2010 (**Fig 1**). Julie Thomas took this strain with her to Rochester Institute of Technology in 2014 (**Fig 1**).

R.A White III purchased the ATCC strain (ATCC_WT) in 2020 but, through cultivation, noticed a larger plaque morphology (RAW_WT;**Fig 1**). Thus, the need to sequence all the strains present in the various laboratories due to spontaneous mutation was warranted due to these changes in phenotype over 50 years, which may be related to single nucleotide polymorphisms, insertions/deletions, and/or methylation status.

Phage G DNA has been anomalous in that (1) pulsed field gel electrophoresis (PFGE) revealed it to be 750 to 630 kbp long (Hutson et al., 1995; Serwer et al., 1995; Serwer & Hayes, 2001) and (2) the NCBI_WT genome size is only 497 kbp by DNA sequencing. Possibly, the difference was caused by a permuted genome with exceptionally long terminal repeated sequences (LTRs). As well, packaging studies have shown that it can package 626 kbp within its capsid head (Ageno et al., 1973; Donelli, 1968; Donelli et al., 1975; 1976; González et al., 2020; Hua et al., 2017). The LTR was not in the NCBI_WT genome draft of phage G. Another possibility is that modifications of G DNA (e.g., methylation, glycosylation, or other unknown modifications) distorted the PFGE. Phage G DNA may also be lethal in *E. coli* upon cloning, making it challenging to resolve LTRs. There is a genome size discrepancy between the DNA packaging, pulse-field gel electrophoresis, the missing LTRs, and observed size from the draft genome (NCBI_WT) genome.

Here, we describe that the phage G genome is not permuted and that the above anomaly is the result of genomic DNA derivatization. In addition, the non-permuted character is unique in that two pairs of otherwise unique DNA ends exist and both pairs have the same terminal repeat. We utilized multiomics to resolve the genome, methylome, and proteome via high throughput Oxford nanopore long read sequencing and particle proteomics Mass Spectrometry via LC-MS/MS. Our study describes in detail the functional repertoire of phage G across five decades of continuous cultivation.

## Results

### Genome features and nearest neighbor comparison

Phage G has a *Myovirus* morphology (i.e., classical T4-like morphology) that is ∼630 nm from head-to-tail (i.e., 450 nm tail and 180 nm head with 5-fold vertices), and it has a genome that is 3x larger than the representative *Myoviridae Escherichia* phage T4 (González et al., 2020; 2021). The phage G genomes RAW_WT, UT_MUT, UT_WT, RIT_WT, and ATCC_WT are ∼499 kbp, have 668-669 protein-coding ORFs, that is AT-rich with a GC content of ∼29% (**Table 1; Fig 2-3**). The annotation predicts that 66% of the genome’s ORFs are hypothetical (i.e., without any known homologs or predicted functions), with a 35 kbp stretch of DNA between positions 291377-327020 containing 66 predicted hypothetical ORFs (**Table S1; Fig 2**). Approximately 40% of phage G genome is split between nucleic acid replication, structural genes (e.g., capsid, tail, portal), and AMGs (**Fig 2-3**)

**Table 1:** Genome Quality Statistics. This table assembly statistics (N90/N50), checkV results which includes MIUViG quality metrics (completeness/contamination), gene count, and GC content (%).

|  | Length | N50/N90 | Provirus | Gene count | Quality | Completeness | Contamination | GC % |
| --- | --- | --- | --- | --- | --- | --- | --- | --- |
| ATCC_WT | 499946 | 499946 | No | 668 | Complete | 100 | 0 | 29.93 |
| NCBI_WT | 497513 | 497513 | No | 663 | High-quality | 100 | 0 | 29.93 |
| RIT_WT | 499819 | 499819 | No | 669 | High-quality | 100 | 0 | 29.93 |
| UT_MUT | 499819 | 499819 | No | 669 | High-quality | 100 | 0 | 29.93 |
| UT_WT | 499819 | 499819 | No | 668 | High-quality | 100 | 0 | 29.93 |
| Moose W30-1 | 419969 | 419969 | No | 542 | Complete | 100 | 0 | 29.72 |

**Figure 2:**
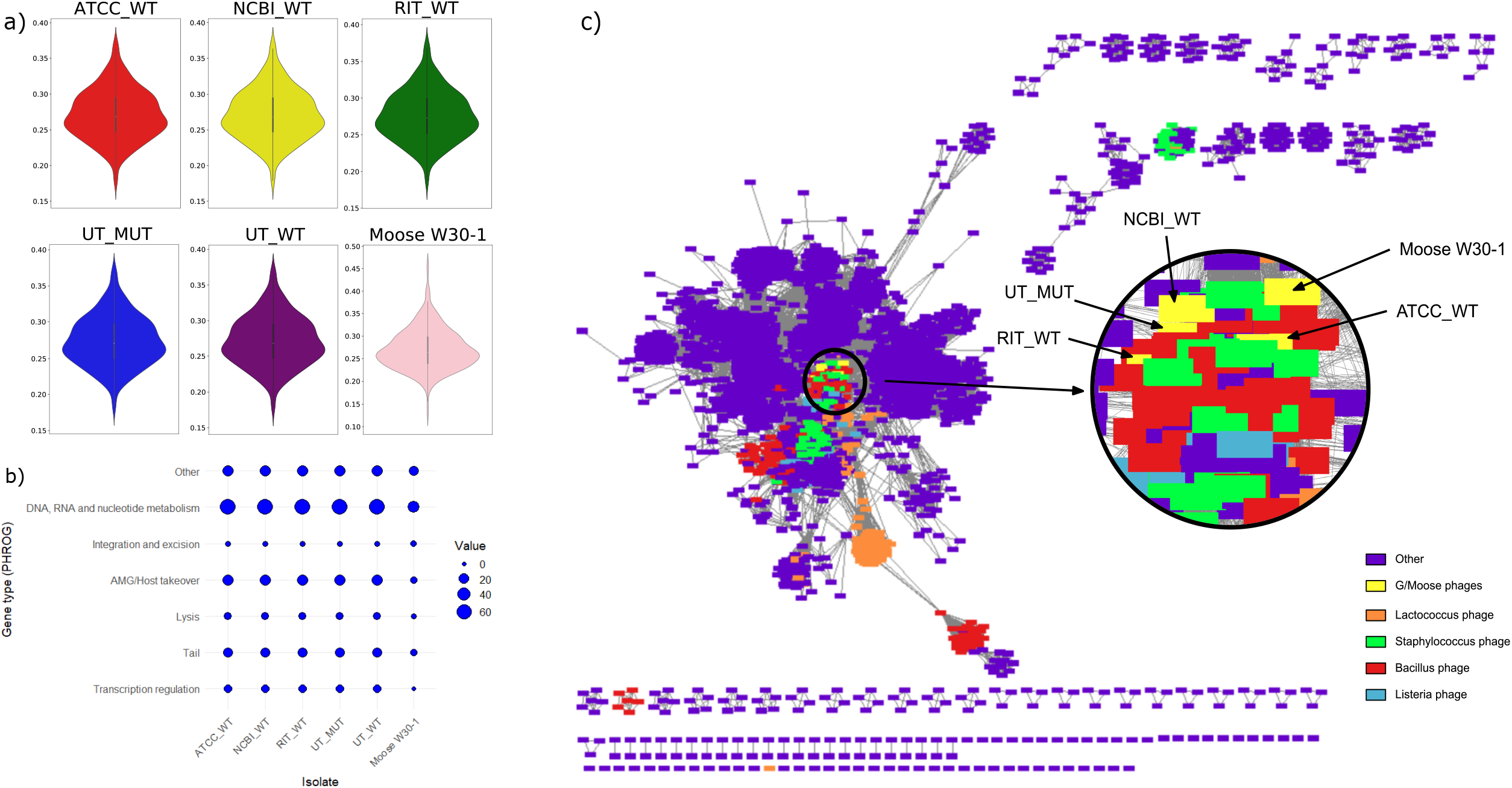
Phage G genomic content and taxonomic placement. A) Violin plots of the GC content across the variants of phage G compared against Moose phage W30-1. B) PHROG hierarchical categories for gene content. Here all phage G gene categories hierarchy are compared to Moose phage W30-1. C) A vContact2.0 plot for taxonomic placement of phage G based on protein/gene content clustering. Moose phage W30-1 failed to be placed.

We applied a network based taxonomic framework to resolve phage G’s relationship to a myriad of uncultivated phages in bacteria and archaea via VConTACT3.0 using INPHARED database (Cook et al., 2021; Bin Jang et al., 2019). All our phage G variants are clustered together with overlap (e.g., RIT_WT, NCBI_WT etc) near each other within a central cluster within our VConTACT3.0 network (**Fig 2**). The phage G cluster also has distant relatives that are *Staphylococcus, Bacillus, Lactococcus*, and *Listeria* phages (**Fig 2**).

PhageAI predicted the following taxonomy and lifestyle for phage G (Tynecki et al., 2020). Phage G has a temperate lifestyle (based on PhageAI at 96.45%), meaning it can enter both lytic and lysogenic lifestyles. The genome has genes whose products are similar to those known to be involved in lysogeny associated integration and excision, including ps548/gp662 transposase (**Table S1**). The genome also has genes for transcriptional regulation (**Table S1; Fig 3a**). The taxonomy is Caudovirales (99.97%) and Myoviridae (97.54%) based on the non-official International Committee on Taxonomy of Viruses (ICTV) taxonomy. TaxMyPHAGE is able to provide the complete ICTV taxonomy, which was recently renamed phage G to *Donellivirus gee* (**Table S1;** Millard et al., 2024). We will use the original name (phage G) for the rest of this paper.

**Figure 3:**
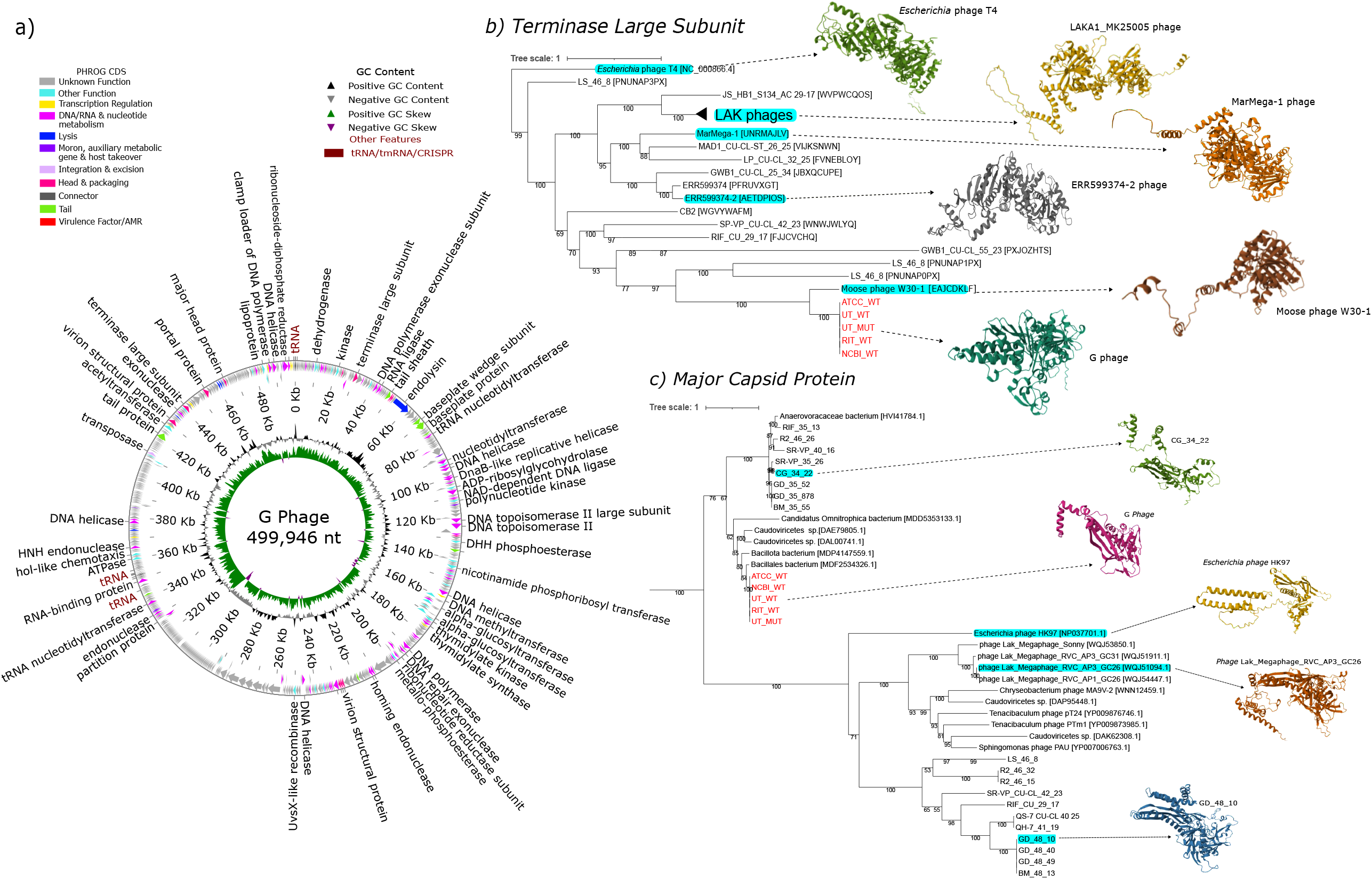
Genomic annotation plot with phylogenetic trees of phage G. A) Genome plot of phage G. This genomic plot of phage G is based on ATCC_WT/RAW_WT. This includes the PHROG CDS annotations, GC content, GC skew, tRNAs/tmRNAs/CRISPRs (phage G has no CRISPRs), and the gene names. B) Large terminase subunit (*TerL*) (ps583/gp1) maximum likelihood tree with 1000 bootstraps.*Escherichia* phage T4 provided the outgroup in this unrooted tree. C) Major Capsid/Head protein (*mcp*) (ps609/gp27) maximum likelihood tree with 1000 bootstraps.*Escherichia* phage HK97 provided the outgroup in this unrooted tree. AlphaFold2 protein predictions are showcased for various representative clades of*TerL*and *mcp*.

The draft genome was published by RWH in 2012. The CheckV result states the NCBI_WT draft genome is high quality (**Table 1**; Nayfach et al., 2021). However, it is incomplete due to the missing LTRs (**Table 1**). The NCBI_WT draft was also missing ∼2 kbp at the 3’ terminal end of the genome when directly compared to RAW_WT, UT_MUT, UT_WT, RIT_WT, and ATCC_WT (**Fig 4)**. All phage G genomes were 99.9% similar to each other based on average nucleotide identity (ANI) (**Fig 4A**). All the genomes (i.e., RAW_WT, UT_MUT, UT_WT, RIT_WT, and ATCC_WT) are at least 499 kbp, suggesting NCBI_WT from RWH either lost ∼2 kbp or was missed in genome sequencing (**Table 1; Fig 4B**).

**Figure 4:**
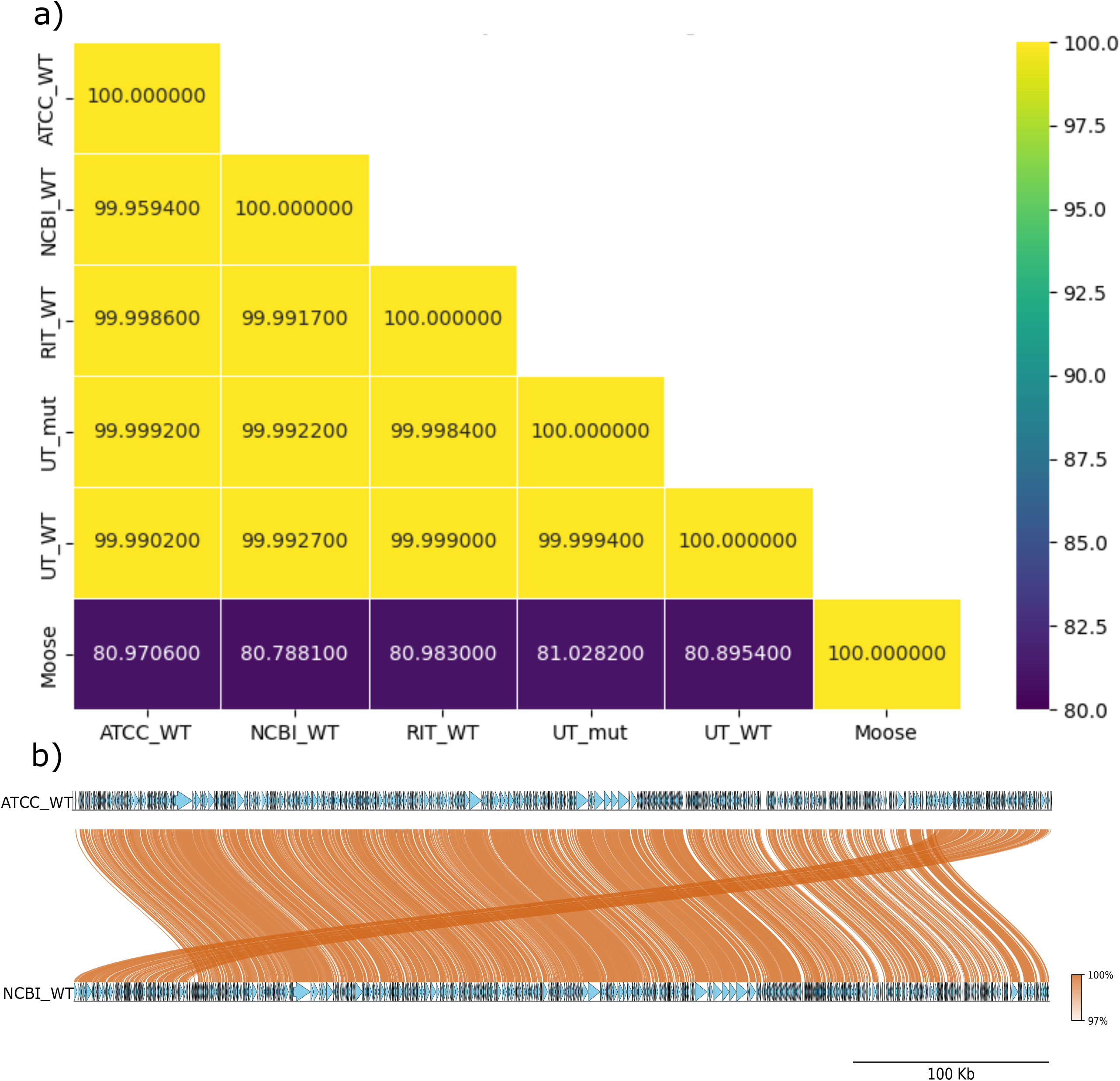
Whole-genome comparison of phage G variants vs. Moose phage W30-1. a) This is an average nucleotide identity (ANI) plot compared using fastANI. b) This whole genome comparison plot of NCBI_WT vs. ATCC_WT.

We performed whole genome alignment using MUMmer (Marçais et al., 2018) and found conserved single nucleotide polymorphisms (SNPs) across all phage G genomes. The conserved SNPs include positions 284289 - G/A, 394072 - T/A, and 395353 - G/A, which are within hypothetical ORFs (ps330/gp428 and ps512/gp625) and a non-coding region (position 394072) (**Table S1**). RIT_WT had more SNPs generally than the other compared variants, including within ps162/gp260, ps206/gp304, multiple in ps485/gp598, within a non-coding region 394072, and within ps578/gp690 (**Table S1**). Most SNPs occur in ps485/gp598, a hypothetical region within RIT_WT (**Table S1**). ORF ps485/gp598 is a hypothetical protein with an unknown functional role (**Table S1**).

In addition, the NCBI_WT genome was artificially rearranged to make the large terminase subunit (*terL*) the starting gene at gp1; which is not the physical starting point of the genome. All five of our variants from independent *de novo* assemblies and long read nanopore data (**Fig 3B**) showed the artificial rearrangement, and that the genome is not a circle (**Table S1**). Thus, we have re-numbered genes based on the actual physical location of genes on the genome as ORFs as ps1-668 (**Table S1**). We used SWORD-based local alignments of the NCBI_WT reference-called ORFs to assign the original gp numbers from NCBI_WT to our annotation (**Table S1**, Vaser et al., 2016). For example,*the terL* gene is at position 440572-442254 and is now ps583/gp1, which maintains the actual physical location plus the original gp number to avoid confusion with previous works over 50 years (**Table S1**).

In addition, we resolved the LTR elements within phage G, due to the use of ultra-long read sequencing with Oxford nanopore.The LTRs are 127 bp in length, with a GC content percentage of 43.31%, which is 13.38% higher GC than the rest of the genome (29.93%) (**Figure 2-3; Fig S1)**. The 5’ and 3’ LTRs are identical, forming terminal genome redundancy. We compared all five genomes (RAW_WT, UT_MUT, UT_WT, RIT_WT, and ATCC_WT), showing the identity of these residues across the LTRs (**Fig S1)**.

We compared our phage G genomes to fifty megaphages present in public databases, including the marine Mar_Mega_1 (Michniewski et al., 2021,**Table S1**). Amongst the genomes listed in ggkbase and Michniewski et al. was a phage genome labeled Moose phage W30-1 (http://ggkbase.berkeley.edu/organisms/405141). Moose phage W30-1 was isolated from moose rumen on a 0.2 μm filter based on ggkbase metadata. While not quite a megaphage of 500 kbp, at only 419 kbp, Moose phage W30-1 is the closest environmental relative of phage G (**Fig 4A**). Moose phage W30-1 is 81% similar based on ANI to phage G (**Fig 4A**). No other phage, whether megaphage or not, was higher than 40% ANI to phage G (**Table S1**). Moose phage W30-1 also clustered with phage G using vConTACT3.0 (**Fig 2**). Moose phage W30-1 has a slightly lower GC percentage than G phage 29.72 vs. 29.93%, it also has 80 kbp less DNA, it has genes with higher GC content up to 49%, and it only has one 5’ LTR (**Table 2; Fig 2**). The LTR present on the 5’ of the Moose phage genome is 99 bp in length and has a GC content of 35.35%; it is not repeated at the 3’ end, suggesting that Moose phage W30-1 has a circular genome, not a linear like phage G (**Fig S2)**.The logo plot suggests a small amount of conservation in the LTRs for phage G and Moose phage W30-1 (**Fig S2**). PhageAI suggested that phage moose W30-1 is a temperate phage just like phage G but at a lower percentage of 60.28%, likely due to containing a similar transposase. Also, like phage G, it is based on non-ICTV taxonomy from phage AI, which is a Caudovirales Order at 99.91% and Myoviridae Family at 97.57%.

**Table 2:** Gene open-reading frames with ≥40% GC content. Both phage G and Moose phage W30-1 have an average of GC content <30%; however, both encode three genes each that have ≥40% GC content.

| ***Moose W30-1*** |  |  |
| --- | --- | --- |
| **ORF** | **Annotation** | **GC%** |
| ORF357 | Hypothetical | 0.49 |
| ORF366 | Hypothetical | 0.48 |
| ORF429 | Hypothetical | 0.47 |
| ***Phage G*** |  |  |
| **ORF** | **Annotation** | **GC%** |
| ps22 | T4-like spike protein | 0.40 |
| ps98 | Tail structural protein | 0.41 |
| ps608 | Capsid decoration | 0.40 |

Despite the >80% ANI between phage G and phage Moose W30-1, both had many unique viral genes. Notable genes found in Moose phage W30-1, but not in phage G, are genes for reverse transcriptase,*HicB*-like antitoxin, tellurite resistance, cytidylytransferase, dCTP deaminase,*NinI*-like serine-threonine phosphatase,*NrdD*-like anaerobic ribonucleotide reductase,*PnuC*-like nicotinamide transporter, and a *RecJ* endonuclease (**Fig 5A**). Comparison by use of MetaCerberus indicated that moose phage W30-1 was relatively enriched for bacterial motility proteins, flagellar assembly, focal adhesion, glycine/serine/threonine metabolism, glycosaminoglycan binding proteins, methane metabolism, and translation factors (**Fig 5B**, Figueroa III, et al., 2024).Pathways of note in phage G include antibiotic biosynthesis, glutathione metabolism, selenocompound metabolism, biofilm formation, and antimicrobial resistance genes (e.g., Dihydrofolate reductase -*DHFR*) (**Fig 5**). The biofilm formation pathway enrichment in phage G relates to spore biosynthesis manipulation, including spore protease (**Fig 5**).

**Figure 5:**
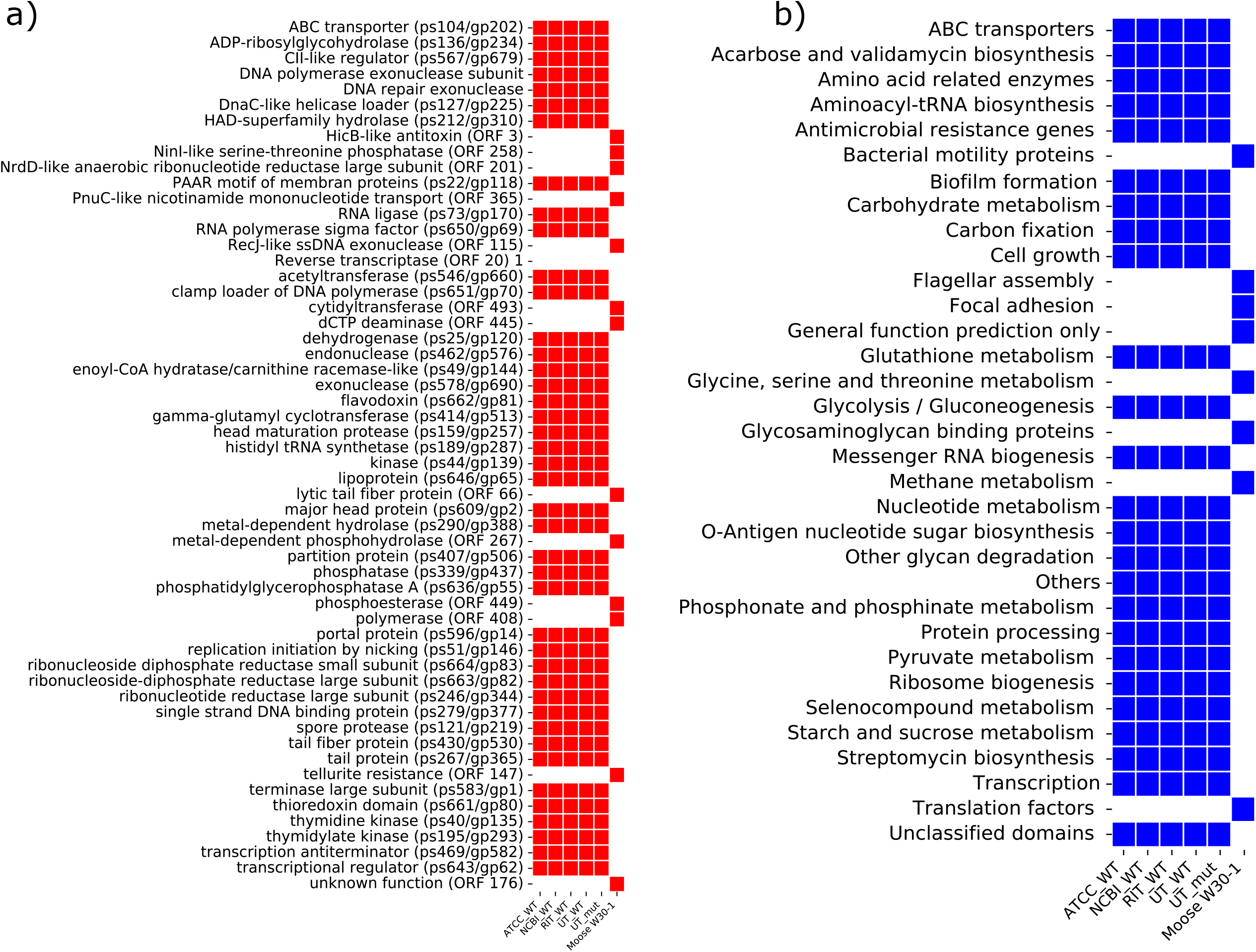
Heatmaps of the most differential genes and pathways comparing phage G variants vs. Moose phage W30-1. a) Annotations of genes b) KEGG pathways. Only genes that are missing from at least one phage are shown.

### Structural genes

Phage G has the hallmark genes of classical tailed phages, including capsid/head, portal, baseplate, terminase (*terL)*, and other structural virion proteins (**Fig 2-3**). Of the many structural genes we detected, sixteen were directly detected with proteomics, which includes the hallmark genes mentioned above (**Fig 6**). Phylogenetic analysis of *the terL* gene suggests that Moose phage W30-1 diverged earlier than all G phage variants (**Fig 3B**). The *terL* of moose phage W30-1 has an extra loop based on the predicted protein structure, which is not found on phage G (**Fig 3B**). LAK phages, phage T4 (i.e., the outgroup), MarMega-1, and ERR599374 phages occur on different branches when compared to phage G and have predicted *terL* structural differences (**Fig 3B**). We could not detect the *terL* protein (i.e., ps583/gp1) within our purified phage proteomics but has been previously detected with proteomics and cryoEM (**Fig 6**, González et al., 2020).

**Figure 6:**
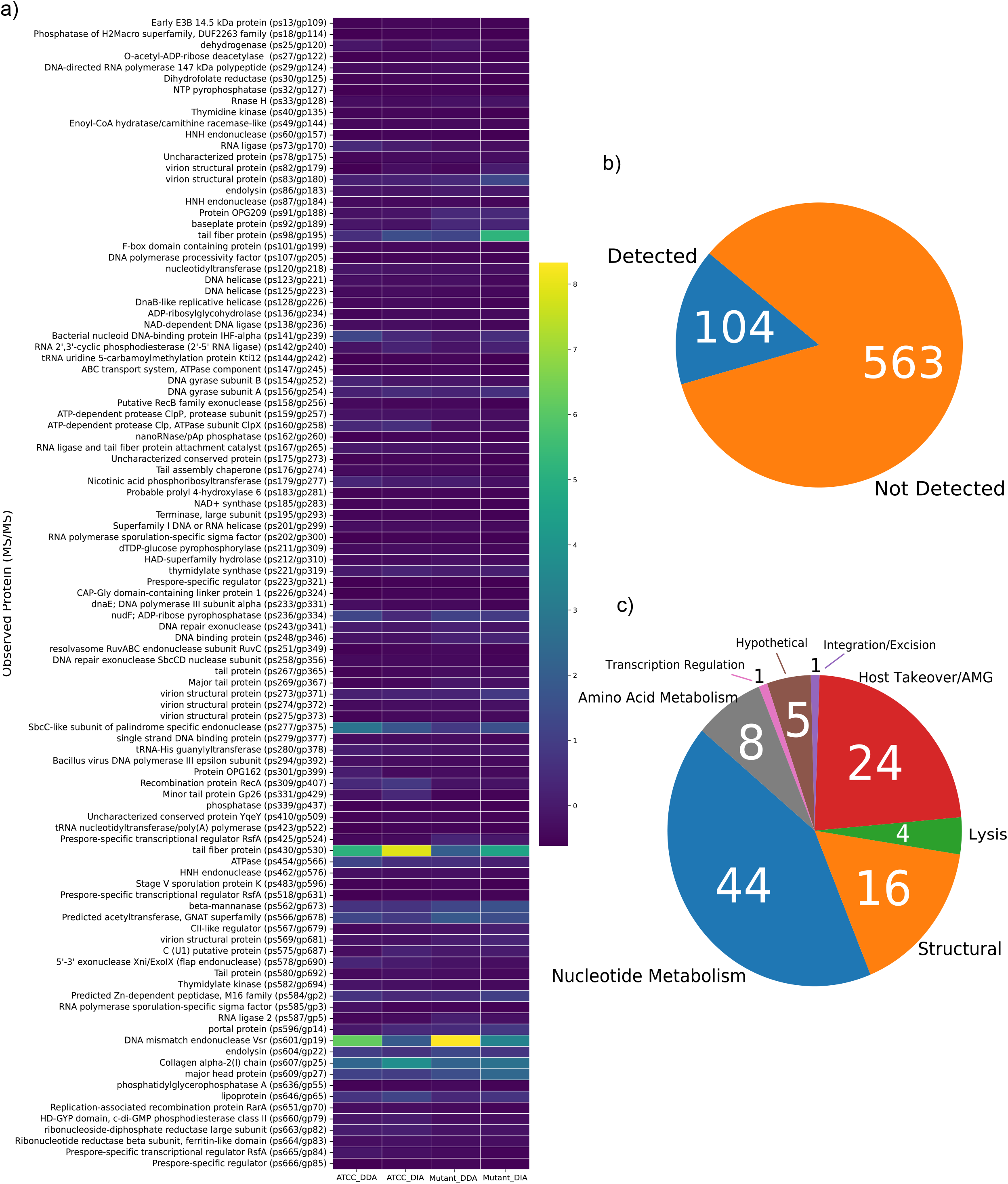
Particle proteomics using LC-MS/MS. a) Heatmap comparing UT_MUT vs. ATCC_WT using DDA and DIA using the log2 normalized values. b) Coverage of the ORFs detected in either UT_MUT and ATCC_WT using DDA and DIA. c) PHROG gene pathways hierarchies in either UT_MUT and ATCC_WT using DDA and DIA

**Figure 7:**
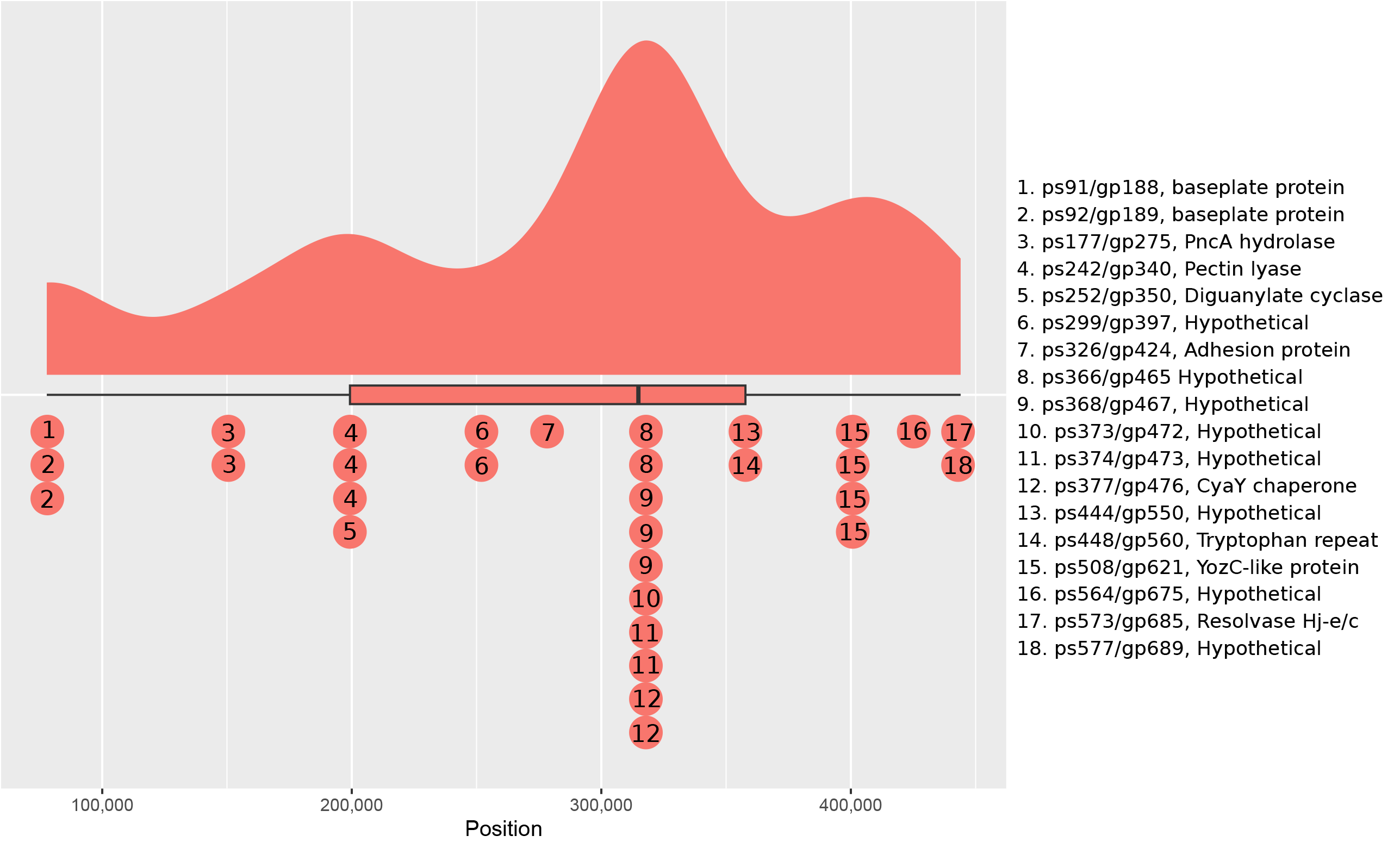
*De novo* methylation call results and positions across the phage G genome plot. To save space the full annotation name was shortened to fit within the plot. This includes the full annotation name for each ORF (ps/gp identification number) is present in Table 3, but for further details check Table S1 annotation for these.

We confirm González et al.’s (2020) results with phylogenetics that place the major capsid protein (*mcp*) (i.e., head with 5 fold vertices) within the HK97 capsid family (**Fig 3C**). The phylogenetics of *mcp* suggests that phage G’s HK97-like capsid diverged earlier than *E. coli* HK97 and is an outgroup compared to it directly (**Fig 3C**). Lak phages and other phages found metagenomically (e.g., GD phages) are also later HK97-style capsids, with *E. coli* HK97 providing an outgroup for them (**Fig 3C**). We have been unable to find the *mcp* in Moose W30-1 phage; it appears highly divergent compared to known *mcp* HMMs (**Table S1, Fig 5A**). We also detect ps609/gp27, the *mcp*, within our proteomic data of purified phage particles (**Fig 5**). A novel capsid decoration protein ps608/gp27 wasn’t detected in our proteomic data but has been previously detected with proteomics and cryoEM (González et al., 2020). ORFs encoding head maturation proteases ps159-160/gp257-258 and ps601/gp19 encoded by phage G were proteomically detected (**Table S1, Fig 6**).

Portal and tail proteins are critical components of every tailed phage particle. The portal protein is encoded by ps596/gp14, which has also been previously detected by proteomics and cryoEM (González et al., 2020); we have also found the protein within our proteomic data both in the ATCC/RAW_WT and the MUT_UT (**Fig 6**). The portal is, based on HHpred and MetaCerberus, related to lambda phage-like portal proteins (pdb:8k39,**Table S1**). Tail and tail assembly proteins include fifteen annotated ORFs, and seven proteins were found by LC-MS/MS for the complex tail assembly in phage G (**Fig 6**). Starting with ps80-86/gp178-183, which includes a head-tail connector similar to SPP1 gp17, contractile tail sheath, the two tail tube subunits 1 and 2, a tail assembly chaperone proteins, programmed frameshift tail chaperone protein, and tape measure protein with tail lysin domain (**Table S1**). Our proteomic data confirms that ps82-83/gp179-180 tail tube subunit 1 and subunit 2 are expressed (**Table S1; Fig 6**). Of the related tail assembly proteins ps98/gp195, ps167/gp265, ps267/gp365, ps430/gp530, ps562/gp673 and Tail fiber protein ps430/gp530 were found within the phage particles’ proteomics (**Table S1; Fig 6**). The tail accessory proteins that were found amongst the LC-MS/MS include a novel CAZy-related GH113 Beta-mannanase lysis protein (i.e., ps562/gp673) that is attached to a tail fiber and ps167/gp265, an RNA ligase with tail fiber protein attachment catalyst (**Table S1; Fig 6**).

The baseplate proteins connecting to the tail include ps91-93/gp188-190, similar to the *P2gpJ/I*-like phage and T4 gp8-like (**Table S1**). The proteomics confirm that ps91-92/gp188-189 that the P2gpJ/I like phage baseplates are expressed; however, ps93/gp190, the T4 gp8-like baseplate was not found within the proteomics (**Fig 6**). HHpred suggested models of Mu gp47/gp48, T4 gp6, and P2 *gpJ/P2gpL* for ps91/gp188 and ps92/gp189 with further matching via HMMs to VOG33143 for ps91/gp188 (**Table S1**).

### DNA replication, repair, and transcription

Genes for nucleic acid metabolism (e.g., endonucleases, exonucleases, and helicases) include sixty-nine annotated ORFs of which 55% are found, expressed, and present within the phage particle by proteomic analysis (**Table S1; Fig 6**). Phage G encodes four DNA polymerases that are DNA pol III-like with two (ps233/gp331 and ps294/gp392) that were within the particle proteomics (**Table S1; Fig 6**). Sixteen endonucleases are encoded by phage G: with the particle proteomics detecting ps60, ps251, ps277, ps462, ps578, and ps604 (**Table S1; Fig 6**). Five exonucleases for DNA repair are also found with four being detected by proteomics (**Table S1; Fig 6**). Protein ps578/gp690 is 5’-3’ exonuclease *Xni/ExoIX* (flap endonuclease) with dual exo-endo putatively, and ps158/gp256 is a *RecB*-like homolog (**Table S1; Fig 6**).

Helicases and helicase-associated ORFs (8 ps/gps), including primase (1 ps/gp), and gyrase subunits A and B (2 subunits), are associated DNA replication with ps154/ps156 (*gyrase A/B*) and ps123/ps125/ps128/ps201 (Helicases) present in the proteomics data (**Table S1; Fig 6**). The helicases ps127-129 are *DnaB/C*-like replicative helicases with loaders, and ps123/gp221 is UvsW-like but not expressed in the proteomics (**Table S1; Fig 6**). Other helicases are viral in nature but are not in known host-related classification families (**Table S1; Fig 6**).

Phage G encodes a gyrase AB subunits (ps154/ps156), and a *UvsX*-like recombinase (*RecA*-like) protein (i.e., ps309/gp407), ribonucleotide reductase (RNR) class 1 b with alpha and beta subunits and *Nrdl*-like flavoprotein-like regulator protein on ps662-664/gp81-83, with ps82-83 detected by viral particle proteomics (**Table S1; Fig 6**). Our phylogenetic analysis of gyrase A and B subunits (ps154/ps156) shows limited relationship to bacterial gyrases, both Moose phage W30-1 and G phage being outgroups within our maximum likelihood estimations (**Fig S2**). Structurally, the *gyrA* subunits between Moose phage W30-1 and G phage are very similar, whereas *gyrB* varies more based on AlphaFold2 (Jumper et al., 2021;**Fig S2**). Our *UvsX*-like recombinase (*RecA*-like) protein based on phylogenetics appears more bacterial based on our maximum likelihood estimations and MAFFT alignments (**Fig S2**). The Moose phage W30-1 *UvsX*-like recombinase (*RecA*-like) outgroups the rest of the G phage, suggesting its *UvsX*-like gene is a more derived ancestor to phage G (**Fig S2**).*UvsX*-like in phage G appears to be horizontally transferred from bacteria based on maximum likelihood, whereas the signal from *gyrA* and *gyrB* is not as clear (**Fig S2**). RNR has essential roles in DNA replication in synthesizing deoxyribonucleotide triphosphates (dNTPs) from ribonucleotides triphosphates (Torrents, 2014). Encoding its own RNR and packaging it within the phage G particle ensures that viral dNTPs are made.

DNA binding proteins for general DNA repair, stabilization, and replication are components of the phage G genome. The genome encodes a thymidine kinase, two thymidylate kinases, and thymidylate synthase (**Table S1**).Thymidine kinase (ps40/gp135) and the thymidylate synthase (ps221/gp319) were detected within the proteomic data (**Table S1; Fig 6**). Three unique DNA binding proteins were found: ps141/gp239, ps248/gp346, and ps279/gp377 with all detected by viral particle proteomics (**Table S1; Fig 6**). In addition, ps141/gp239 encodes a bacterial nucleoid DNA-binding protein IHF-alpha, which is histone-like (Travers, 1997,**Table S1; Fig 6**). ORF ps248/gp346 encodes a ferritin-like DNA-binding protein often involved in DNA replication (Smith, 2004,**Table S1; Fig 6**). Gene product ps279/gp377 encodes a single-stranded DNA-binding protein usually involved in DNA replication (Maffeo & Aksimentiev, 2017). Protein ps251/gp349 contains holliday junction resolvase *RuvABC* endonuclease, also detected in proteomics data (**Table S1; Fig 6**).

We predicted the origin of replication (*OriC*) and various promoter motifs across the G phage genome with two putative *Ori*Cs and 2852 putative promoters (**Fig S3; Table S1**). The first predicted OriC starts 1-406 nucleotide position within the genome, has 52% A+T content, and has the classical 5’/3’*DnaA* boxes, spacer,*DnaA* trio, ATP-DNA box, AT-rich region,*CtrA* and Fis binding sites/motifs, and IHF binding site (**Fig S3**). The second predicted *OriC* is ∼200 bp upstream of the first predicted *OriC* at position 611-986 bp and has all the classical site/modify predictions as the first, with a higher A+T content of 58% (**Fig S3**). Of the predicted promoters, 97.5% were host promoters, and <3% were predicted to be phage promoters (**Table S1**). All predicted promoters had a >50% score, with 975 having a >85% predicted score, 720 having 90% or above, and 60 at 100% (**Table S1**).

While host-encoded promoters dominate the predicted promoter landscape of phage G’s genome, we focused on high-quality phage-related promoters. We predicted 19 at 80% or above and nine at 90% or above, with more GC content than the host-related putative promoters (**Table S1**). The phage promoters with the highest scores rounded to 90% or above are promoters for multiple non-coding regions or are within coding regions within phage G. We have nine predicted phage promoters with 222643-222665 and 139516-139538 having the highest predicted scores 98.4% and 97.2% (**Table S1**). Promoter on position 222643-222665 is a short motif ‘TAATACTACTCACTATATCAGAG,’ within ps269/gp367 is a hypothetical ORF with no functional annotation. The ORF ps269/gp367 is a *Bacillus* conserved hypothetical ORF related to phrog_11333 (6.1E-178) that is often co-localized with phrog_16766 a host chromosome condensation regulator (**Table S1**). Phage promoter at 139516-139538, a short motif ‘AAATAACCTTCAATAAGAGGATA,’ within a hypothetical protein (ps172/gp270).

Position 136897-140626 has seven predicted ORFs with no known function; only ps167/gp265, an upstream RNA ligase and tail fiber protein attachment catalyst, and ps175/gp273 downstream MutT/NUDIX hydrolase have predicted function (**Table S1**). While it is a highly confident promoter prediction, its regulation of function or function itself is unknown.

Transcription and host-related transcriptional regulation are highly present and adaptive within phage G’s genomic repertoire. Phage G encodes many host transcription factors, including four *RsfA* family-like, three RNA polymerase sigma-70 like factors, Mu-like gp9 factor, and three factors that are listed prespore or unlisted protein family relationship (ps223/gp321, ps281/gp379, ps643/gp62) (**Table S1, Fig 6**). Phage G encodes an antiterminator (i.e., ps469/gp582), which regulates host RNA transcription in other phage models (Conant et al., 2008). These antiterminators are often co-located with terminase small subunit (*TerS*), which we could not find.

### Translation and lysis

As with nucleic acid metabolism and replication, phage G controls its translation, including its regulation within the host cell. Phage G encodes its own tRNAs and has 18 of them, with 1 tmRNA predicted (**Table S1**). The tmRNA is encoded in position 352812-353120, with a target peptide of ‘AKLNITNNELQVA*,’ within a non-coding region between ps444-ps445/gp550-gp556 and is 308 bp in length (**Table S1**). The ps444-ps445/gp550-gp551 regions are of unknown function, but ps445/gp556 is predicted to be a tryptophan repeat gene family of unknown function. The tRNA gene encoding occurs at the near beginning of the genome position 48-1431, which encodes 6 tRNAs, and towards the end of the genome position 344693-499942 (**Table S1; Fig 3**). The tRNAs encoded by the genome include Arg: TCT, Asn:GTT, Asp: GTC, Cys: GCA, Gln:TTG, Gly: T.C.C., His:GTG, Met: CAT, Thr: TGT, Trp: CCA, and Tyr: GTA (**Table S1; Fig 3**). It has tRNAs with multiple copies, including Glu: TCC with two copies, Phe: GAA with two copies, and multiple alternative tRNAs for serine (Ser:TGA and Ser: GCT) (**Table S1; Fig 3**).

There were eight detected translation/amino acid metabolism-related genes found in the viral particle mass spectrometry data, including *ClpP* (ps159/gp257),*ClpX* (ps160/gp258), prolyl-4-hydroxylase 6 (ps183/gp281), and phosphatase (ps339/gp437) (**Table S1; Fig 6**). The genome encodes five individual proteases, including *ClpP* and *ClpX*, a head/capsid maturation protease (ps19/gp601), membrane protease (ps641/gp60, a *YdiL*-like, which is within the CAAX protease family), and a spore protease (ps121/gp219). Only *ClpP* and *ClpX* were detected in the proteomic data; none of the other proteases encoded were detected in the proteomic data (**Table S1; Fig 6**).

All phages that can enter a lytic phase of their life cycle, whether temperate or strictly lytic, must escape their host cell via lysis of the cell membrane. Phage G has six genes with known homology to endolysins, holins, and spanins. These lysis genes are encoded near the end of the genome from position 373683-463039, encoding the three endolysins, the holin, and an *Rz*-like spanin protein (**Table S1**). ORFs ps604-605/gp22-23 encode similar N-acetylmuramoyl-L-amidase/murein transglycoslyase/flgJ muramidase (EC 3.5.1.28) endolysins with ps604/gp22 but not ps605/gp23 detected within the proteomic data (**Table S1; Fig 6**). ORF ps575/gp687, which encodes the Rz-like spanin, was also detected in the proteomic data (**Table S1; Fig 6**). ORF ps475/gp588 encodes a VOG62162 and phrog_25820 endolysin; however, it has homology to KO number K23989, which is a Mannosyl-glycoprotein endo-beta-N-acetylglucosaminidase which has *lytD* and *lytB* domains but is not found within the proteomic data (**Table S1; Fig 6**). ORF ps86/gp183 is a novel tape measure and tail assembly chaperone protein with an attached tail lysin. This protein was detected in the proteomic data and may be a Tailocin-like endolysin (**Table S1; Fig 6**). A holin protein is also encoded by ps607/gp25, which was detected in the proteomic data, and was previously found in prior proteomics (González et al., 2020).

### Host response and Auxiliary Metabolic Genes (AMGs)

Phage G has two genes that appear to directly interfere with host antiviral defense: anti-CBass nuclease *Acb1* (ps20/gp116) and Anti-Pycsar protein *Apyc1* (ps290/gp388) (**Table S1**) (Hobbs et al., 2022). These proteins are part of the cyclic oligonucleotide-based antiphage signaling system (CBASS) and the pyrimidine cyclase system for antiphage resistance (Pycsar) that resist host antiviral immune response (Hobbs et al., 2022). The closest hits for ps20/gp116 are to pVOG7183 (Unannotated protein) and phrog_12204 (unknown function), with phrog_12204 often co-localized with phrog_22915 a membrane protein involved in the moron, AMG and host takeover category. HHpred matches ps20/gp116 to pdb:7T26_A Anti-CBass nuclease *Acb1* and Uniprot-sprot-vir model P04533 (T4 gene 57B). ORF ps290/gp388 has homology to VOG01700 which is Anti-Pycsar protein *Apyc1* (5.4E-67,**Table S1**). Neither ps20/gp116 nor ps290/gp388 were found amongst the proteomic data (**Table S1; Fig 6**).

Dihydrofolate reductase ( *DHFR*) and *PhoH*-like genes are encoded by phage G. Phage G, on ORF ps30/gp125, has a homolog to a *DHFR* as first described in Enterobacteria phage T4. We applied phylogenetics to phage *DHFR*s with the *Enterobacteria* clade containing T4 with relatives AR1 and *Serratia* phage Muldoon forming the outgroup. Phage G forms a unique clade that branches from *Citrobacter* phage Mijalis and Maleficent and mainly contains gram-positive *Bacillus* and *Salimicrobium* (**Fig S4**;**Table S1**). Moose phage W30-1 also has *DHFR*,which shares some structural homology based on alphafold to phage G; however, as with other related phylogenies where Moose phage W30-1 has a homolog with phage G, it appears to have diverged earlier (**Fig S4**;**Table S1**). Phage G’s *phoH*-like protein is encoded by ps65/gp162 and ps189 in Moose phage W30-1 (**Table S1**). ORF ps65/gp162 was not found proteomically in the virion particle (**Table S1; Fig 6**). We compared *phoH’*s from gram-positive relatives and then used *Escherichia* phage T5 as an outgroup; we found again that the Moose phage W30-1 diverged earlier and is in a separate clade when compared to phage G *phoH*, which forms a unique clade that further outgroups from a *Bacillales* bacterium (**Fig S4**;**Table S1**). The phage G and Moose phage W30-1 predicted structures are highly divergent, and the phage G protein is more complex than *Escherichia* phage T5 *phoH* but less complex than Moose phage W30-1 (**Fig S4**;**Table S1**).

Phage G has proteins related to flagellar operon protein (TIGR02530) (ps41/gp136), a *FtsZ*/tubulin-like GTPase (ps43/gp138), and F-like type IV secretion system proteins (T4SS) homologs (ps58-59/gp155-156) (**Table S1; Fig 6**). The ps41/gp136 and ps43/gp138 were not detected proteomically (**Table S1; Fig 6**). The ORFs ps41/gp136 within the TIGR02530 family are located between genes *flgD* and *flgE* (Mukherjee & Kearns, 2014). ORF ps43/gp138 is a *FtsZ*/tubulin-like GTPase, which also has a homolog to KEGG KO *CetZ* K22222 (**Table S1**).

*Pseudomonas* phages provide an outgroup and structurally similar folds to phage G but are still highly diverged phylogenetically and structurally (**Fig S5**;**Table S1**). The phage G version appears more bacterial than phage-like as it forms a unique clade with *Thermosipho* sp as an outgroup (**Fig 5**;**Table S1**). Phage G has two ORFs predicted to be *TraC*-like and *TraD*-like homologs within the T4SS secretory system ps58-59/gp155-156, which were not expressed in the virion particle proteomics (**Table S1; Fig 6**).

Phage G possesses a multitude of sporulation genes to manipulate its *Bacillus* host sporulation. These sporulation related genes include a spore protease (ps121/gp219), small acid-soluble spore protein D (minor alpha/beta-type SASP) (ps168/gp266), RNA polymerase sporulation-specific sigma factor (ps202/gp300), prespore-specific regulator (ps223/gp321), Prespore specific transcription factor without *RsfA*-like domain (ps281/gp379), *YtxC*-like sporulation protein (ps306/gp404), stage V sporulation protein K (AAA-like ATPase, ps483/gp596), and three prespore-specific transcriptional regulators RsfA-like (ps518/gp631, ps665/gp84, ps666/gp85) (**Table S1**). Of the sporulation related genes in phage G, only ps223/gp321, ps483/gp596, ps518/gp631, ps665/gp84, and ps666/gp85 were expressed in the virion particle proteomic data (**Table S1; Fig 6**). Moose phage W30-1 had only one spore-related gene, ps119, which is similar to phage G’s RNA polymerase sporulation-specific sigma factor (ps202/gp300) (**Table S1; Fig 6**). We further resolved stage V sporulation protein K (ps483/gp596) using phylogenetics, which is a K06413 homolog (gene ID *spoVK*) that also matches VOG44220 (AAA family ATPase). The closest clade to phage G’s spoVK was *Clostridium botulinum* related (**Fig S4**). Phage Moose W30-1 did not have a homolog to sporulation protein K (ps483/gp596), nor have we found a phage that does only allowing us to directly compare to spore-forming gram-positives such as *Bacillus* and *Clostridium* (**Fig S4**).

### Methylome

We evaluated the whole-genome methylation landscape of phage G using DeepSignal2 (Ni et al., 2019). We found 31 methylation sites across the genome significantly above the confidence threshold (**Table 3; Fig 6**). Of all the methylation modification of the phage G genome, 71% occurs after position 251905, and ∼40% of the methylation occurs between 314147-363191 (**Table 3; Fig 6**). Approximately 32% of the methylation within the genome occurs in the cryptic region which is a 35 kbp stretch (i.e., positions 291377-327020) that has 66 ORFs that are hypothetical with no known homology but to phage G itself (**Table 3; Fig 6**). Both MetaCerberus, SWORD, and HHpred could not find matches to the various ORFs within this highly methylated region, so we used foldseek to help us resolve these ORFs (Van Kempen et al., 2024). Two ORFs with known annotation upstream and within the unknown zone include the gp424 adhesion protein family that is only found in phage G (i.e., ps326/gp424, methylation position 278258) and ps377/gp476, which is a *CyaY* chaperone-like protein (methylation position 321477 and 321589) (**Table 3; Fig 6**). The ORFs ps366/ps368/ps374 have multiple methylations but have unknown functions as methylation ‘hotspot’ within the genome (**Table 3; Fig 6**). Methylation positions 77050, 77779, and 78199 are within the baseplate protein assembly ORFs ps91-92/gp188-189, where long tail fibers attach to phage particles (**Table 3; Fig 6**). Three ORFs with multiple methylation sites include ps177/gp275, ps242/gp340, and ps508/gp621 which encode a Pyrazinamidase/nicotinamidase/isochromatase hydrolase *PncA*-like protein, a Trimeric auto-transporter adhesins pectin lyase (DUF2807-like), and *YozC*-like protein (**Table 3; Fig 6**).

**Table 3:** *De novo* methylation call results and positions across the phage G genome. This includes the full annotation name for each ORF (ps/gp identification number) but for further details check Table S1 annotation for these.

| ORF | GP | Annotation | Location |
| --- | --- | --- | --- |
| 91 | 188 | P2gpJ-like baseplate protein | 77050 |
| 92 | 189 | P2gpI-like baseplate protein with C-terminal appendage | 77779, 78199 |
| 177 | 275 | Pyrazinamidase/nicotinamidase/isochromatase hydrolase PncA-like | 150529, 150535 |
| 242 | 340 | Trimeric auto-transporter adhesins pectin lyase (DUF2807-like) | 199184, 199227, 199234 |
| 252 | 350 | Probable diguanylate cyclase DgcQ | 199320 |
| 299 | 397 | Hypothetical | 251905, 252007 |
| 326 | 424 | gp424 adhesion protein family (Gphage) | 278258 |
| 366 | 465 | Hypothetical | 314147, 314150 |
| 368 | 467 | Hypothetical | 314848, 314854, 314866 |
| 373 | 472 | Hypothetical | 318043 |
| 374 | 473 | Hypothetical | 318860, 318862 |
| 377 | 476 | CyaY chaperone-like protein | 321477, 321589 |
| 444 | 550 | Hypothetical | 352329 |
| 448 | 560 | Tryptophan repeat gene family protein | 363191 |
| 508 | 621 | YozC-like protein | 400592, 400769, 400860, 400912 |
| 564 | 675 | Hypothetical | 425189 |
| 573 | 685 | Holliday junction resolvase Hj-e/c-like | 441939 |
| 577 | 689 | Hypothetical | 443970 |

### Particle biophysical characteristics

We performed various tests to measure the biophysical stability of the particles under abiotic environmental stressors (i.e., temperature and pH) for wild type vs. mutant (i.e., RAW vs. UT_MUT) phage G. This was to determine if such stressors would impact the infectivity of phage G in the wild type or the UT mutant strain. Each stressor was for 30 mins of time, followed by recovery to the specific pH or temperature. Generally, the UT_MUT has more variance of pfu/mL^-1^ than RAW_WT exempt 40 °C and pH 4, where RAW_WT had more (**Fig 8**). The UT_mut is more stable during temperature increases from 40-60 °C compared to the RAW_WT strain (**Fig 8**). At 40 °C, the UT_MUT has nearly half a log more pfu/mL^-1^ than WT_RAW (**Fig 8**), which was statistically significant (p < 0.05). RAW_WT appeared more sensitive to heat than UT_MUT, but it had a more significant variance of pfu/mL^-1^ (**Fig 8**). We attempted a 70 °C treatment, but no plaques were obtained for either the wild or mutant type.

**Figure 8:**
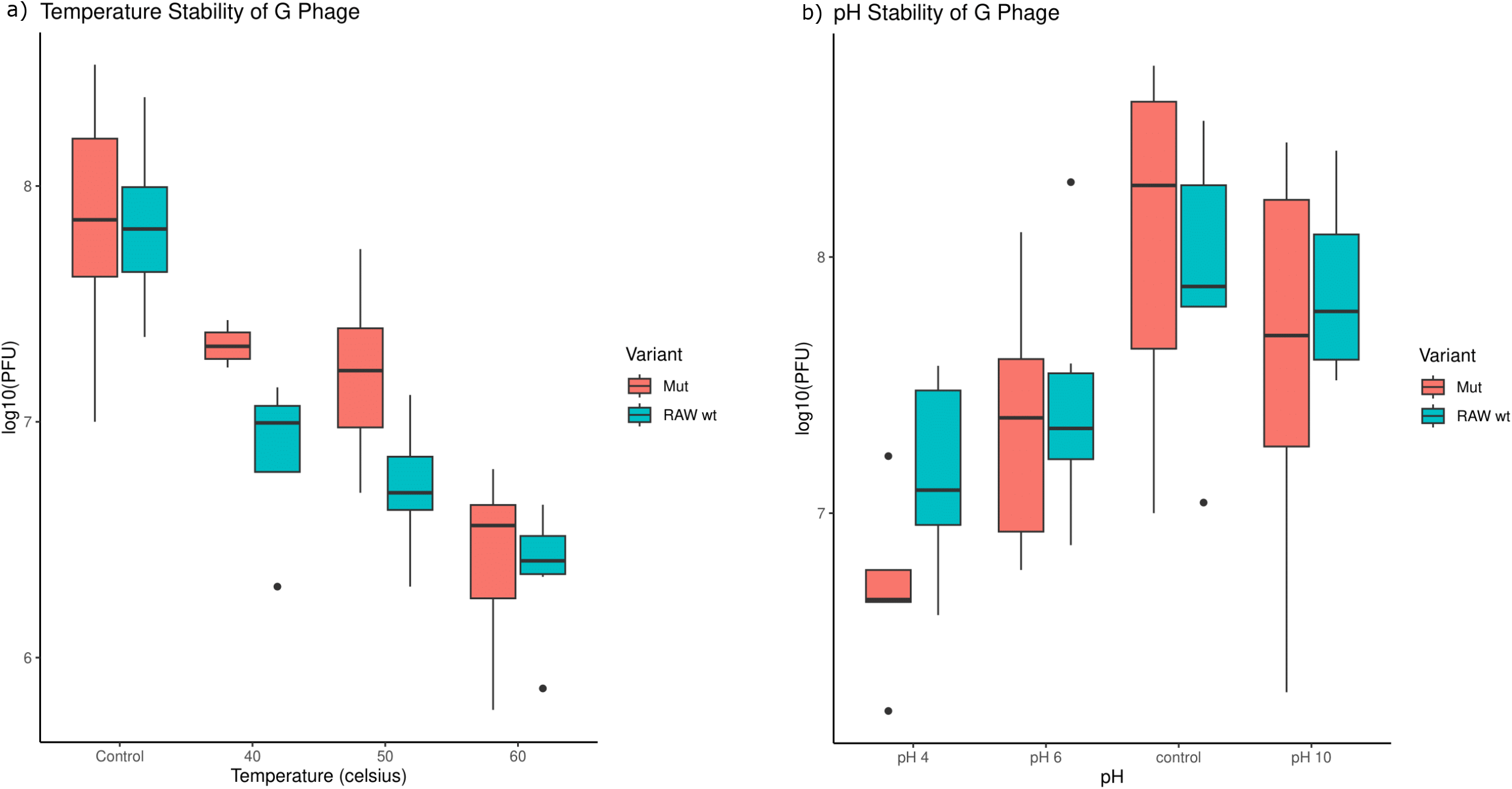
Particle biophysical characteristics of phage G RAW_WT vs. UT_MUT. a) Thermal stability test. b) pH stability test. Each stressor was for 30 mins of time followed by recovery to the specific pH or temperature.

The UT_MUT had more significant variances of plaque formation at pH 6, control pH 7, and alkaline pH 10 (**Fig 8**). UT_MUT lost nearly one entire pfu/mL^-1^ unit at acidic pH (pH = 4), which was statistically significant, but RAW_WT had a higher variance of pfu/mL^-1^ at pH four than UT_MUT (**Fig 8**). We attempted pH 3 and pH 11, but there were too few plaques to count, and they were highly inconsistent in forming plaques amongst replicates; thus, they were omitted.

## Discussion

Here, we have presented the methylome, particle proteome, the biophysical properties including biochemical stability of the phage particle, and robust genomic annotation for the only known cultivated megaphage on Earth, The phage G genome is ∼499 kbp at 29% GC content, containing 668 predicted ORFs, of which 104 are proteomically detected within the phage particle. Comparing the five variants of phage G, they are 99.9% similar with a few conserved SNPs and are similar to Moose phage W30-1 that has never been cultivated.

Based on our data, the sequence discrepancy with the PFGE data is caused by modification of the bases in the phage G genome, not by an unusually long terminal redundancy. Indeed, our data indicates that phage G is highly methylated within a cryptic region (i.e., positions 291377-327020). Methylation can impact the mobility of DNA through gels, as shown by (Kinoshita-Kikuta et al., 2008). Further experimental validation is needed related to methylation with PFGE studies.

PhageAI predicted that phage G was a temperate phage, which means that it can enter both the lysogenic and lytic phases. Only a few genes, such as a transposase, were found to possibly signify lysogenic lifestyle. However, lysogenic lifestyle has never been observed for phage G host in 50 years. CheckV results suggest phage G is not a prophage nor does it exist as a prophage within its host *Lysinibacillus* sp. PGH (González et al., 2020). Lytic/lysogenic classification for phage G is not certain.

Furthermore, we have resolved the genome to completeness with no gaps without contamination, correcting a decades-long debate by finding the missing that the LTR was 121-bp. We also found a missing 2 kbp at the 3’ end of the draft NCBI genome. The genome is linear with ends that require revision of the starting point of the sequence in NCBI_WT. All five variants of G phage were 99.9% the same based on ANI. The NCBI published variant is no longer available. Thus, we could not determine whether the 2 kbp was missing in the NCBI_WT variant. Sanger sequencing of phage G DNA might be challenging because (1) phage G DNA is highly methylated, which would make cloning difficult, and (2) some genes could be lethal in an *E. coli* cloning vector. Either of these two factors could be the cause of the missing 2 kbp piece in the NCBI reference. In addition, LTRs and repetitive sequences are also hard to resolve at the read length of Sanger sequencing (∼600-800 bp).

Our annotation corrected the physical orientation and robustly improved it. Given the above revisions, we have also renamed the ORFs as ‘ps’, the initials (i.e., in honor of Philip Sewer) of the laboratory that kept phage G active (frozen stocks of phage G are unstable) and was the immediate source of the versions sequenced here. We retained the original gp numbering from the NCBI reference to avoid confusion. However, logically, we think that future work on phage G should use the new gene numbering system, which we will update biyearly with the advent of new technology or functional discoveries of the hypothetical ORFs within phage G.

While our study provides a robust annotation, still 66% of the genome ORFs are hypothetical even with modern structural AI-based predictions (e.g., foldseek), HMMs (e.g., MetaCerberus and HHpred), and standard alignment approaches (e.g., SWORD). Furthermore, the role of the cryptic zone (i.e., a 35kbp track of the genome with limited functional annotation where 32% of all genome methylation occurs) is not known. The methylome of phage G provides an excellent opportunity for further discovery. CRISPRi or other genome-level manipulations, including protein-protein interactions, are recommended as future directions to resolve the functions of phage G (Adler et al., 2023).

Our results are confirmatory to various previous proteomics and cryo-EM done previously on the particle capsid/head assembly and other proteins (González et al., 2020). We went further to show that, based on phylogeny, the *mcp* and *terL* phage G classifies in a unique megaphage group that is unlike others found. Besides Moose phage W30-1, these genes appear to be very different from those of the other Lak phages and megaphages like MarMega-1. Phage G’s *mcp* (i.e., HK97 gp5 lineage) is 60% larger than its counterpart in *Escherichia virus* T4 (gp23 lineage) (González et al., 2020). Phage G’s *mcp* derived earlier than HK97 or the Lak megaphages based on our phylogenetic analysis. This would suggest that phage G *mcp* is potentially older or more divergent than Lak or HK97 capsids. The tail assembly of phage G is highly complex, but various lysis proteins are connected to ensure entry and lysis.

Nucleic acid metabolism, repair, and transcription appear highly complex and regulated within G phage, as it is able to replicate 499 kbp of its genome in under 2 hours. Various proteins related to nucleic acids were discovered within the phage particles of phage G, including DNA polymerases, exonucleases/endonucleases, helicases, primases, gyrases, and *UvsX*-like recombinase. Various DNA repair mechanisms exist within phage G, which may account for its genome SNP stability over five decades across multiple labs. We have also identified the potential *OriC* of phage G, which begins at the start of the genome, and have resolved multiple phage/host promoters in need of functional validation. Phage G may also control host transcription via antitermination, which prevents early termination of RNA transcription, and regulates phage lifestyle including initiating lytic or lysogenic phases (Conant et al., 2008; Murchland et al., 2020). Proteomics detected multiple genes within the particle that can potentially protect phage G from DNA damage (e.g., *RecA*-like, histone-like and ferritin-like binding proteins). The resolvase *RuvABC* endonuclease, thymidine kinase, and thymidylate synthase encoded by phage G have yet to be functionally validated. Thymidine and thymidylate-related genes function to repair thymidine dimers (Foekens et al., 2001).

*RuvABC*-like (i.e., ps251/gp349) as host proteins are involved in DNA repair and recombination including resolving cruciform DNA (Fogg et al., 1999; Amit et al., 2004). Phage G (i.e., ps141/gp239) encodes a histone-like DNA-binding protein which can be involved in phage recombination (Travers, 1997). The ferritin-like DNA binding proteins (ps248/gp346, ps279/gp377) that may bind DNA and/or metal ions during DNA replication (Smith, 2004; Maffeo & Aksimentiev, 2017).

Phage G encodes its own translation machinery including many of it is own tRNAs, a suppressor tRNA, various proteases, including *ClpXP* (ps159-160/gp257-258) to monitor translation, and various enzymes to lyse the cell. Phage G encodes a suppressor tRNA or tRNA Sup:UUA that is not associated with the translation of any known amino acid (Santos et al., 2022). It is a UUA anticodon (i.e., RNA Sup:UUA) that decodes the TAA stop codon, which is rarely used in bacteria (0.79%) but more commonly found in eukaryotes (∼21%) (Santos et al., 2022). *ClpP* is an ATP-dependent protease that is involved in head mutation and virion structural formation. *ClpP* and *ClpX* interact to degrade misfolded proteins when the cell is under stressful conditions, allowing cellular protein stability (Krüger et. al., 2000). Spanins encoded by ORF ps575/gp687 (i.e., Rz-like spanin) are more commonly found amongst gram-negatives, not gram-positives, as they are required to disrupt the outer membrane in gram-negatives (Kongari et al., 2018). We confirmed González et al. (2020) finding of the ps607/gp25, a holin protein which is a small protein that punches holes in bacterial cell walls towards the end of the lytic lifestyle towards particle release (Wang et al., 2000).

Phage G genome encodes homologs to flagellar operon protein (TIGR02530) (ps41/gp136), *FtsZ*/tubulin-like GTPase (ps43/gp138), and F-like type IV secretion systems (T4SS) homologs (ps58-59/gp155-156). ORFs ps41/gp136 is a TIGR02530 family gene that is in-between genes *flgD* and *flgE*, which are flagella hook formation (Mukherjee & Kearns, 2014). *FlgE* is the main structural subunit of the flagella hook protein, and *FlgD* polymerizes *FlgE* protein and provides scaffolding for the flagella hook in *Bacillus subtilis* (Mukherjee & Kearns, 2014). In bacteria, *FtsZ* assembles a Z-ring, which is the site of cell division and is essential for the process; it also has roles in peptidoglycan synthesis and is the bacterial homolog for eukaryotic tubulin (Margolin, 2005). Homologs of *FtsZ* or *TubZ* in phages are commonly found in gram-negative bacteria (e.g., *Pseudomonas*) and their phages, including phiKZ, which we used as an outgroup in our phylogenetics (Oliva et al., 2012; Aylett et al., 2013). *FtsZ/TubZ* homologs in phages act as DNA partitioning systems and cytomotive GTPases and may act as transporters of phage DNA to cellular poles (Oliva et al., 2012). F-like type IV secretion systems (T4SS) transport DNA between bacterial cells (Wallden et al., 2010). ORFs ps58-59/gp155-156 are *TraC*-like and *TraD*-like homologs within the T4SS secretory system. *TraC* and *TraD* are membrane-bound ATPase proteins within the bacterial cell, which provide energy in the form of ATP for pilus extension and transferosome transport reciprocally (Bragagnolo et al., 2020). Their function within phages is entirely unknown.

Phage G on ORF ps30/gp125 has a homolog to a *DHFR*. *DHFR* was first described in *Enterobacteria* phage T4 back in 1970; in T4, it functions in building tetrahydrofolate for thymidylate synthesis, which supplies nucleotides for DNA synthesis and has structural roles as a protein in the phage baseplate of the tail (Kozloff et al., 1970; Mosher et al., 1979). The impact of the trimethoprim resistance being conferred or transduced from phage to bacterial host via a *DHFR* has not been functionally studied. Further investigation of *dhf’s* within phages warrants further study.

Another AMG within phage G and in Moose phage W30-1 includes *phoH* -like homology, which is found amongst many phages but represents a first within megaphage. *PhoH* is part of the *Pho* (phosphate) regulon genes, which are induced in phosphate starvation; it is a cytoplasmic protein that is predicted to be an ATPase with ATP binding activity in *E. coli* K-12 (Kim et al., 1993; Koonin & Rudd, 1996; Metcalf et al., 1990). The gene is highly prevalent within phage genomes within marine ecosystems at ∼40%, with ∼4% in non-marine phages (Goldsmith et al., 2011). No phage functional analysis has been completed in a phage *phoH* ; thus, its function still needs to be discovered. Further functional analysis is warranted in *phoH* , generally including the megaphage versions in phage G and Moose W30-1.

Phage G has antiviral escape genes anti-CBass nuclease *Acb1* (ps20/gp116) and Anti-Pycsar protein *Apyc1* (ps290/gp388), which may allow it to escape host response. Anti-CBass nuclease *Acb1* hydrolyzes a tricyclic nucleotide used by the host to signal cell death and host immunity in response to phage infection (Hobbs et al., 2022). The first *Apyc1* protein was functionally validated in *Bacillus subtilis* phage SBSphiJ as a metal-dependent cyclic NMP phosphodiesterase that targets pyrimidines (mainly cytidine 3’,5’-cyclic monophosphate (cCMP) and uridine 3’,5’-cyclic monophosphate (cUMP) (Hobbs et al., 2022). Both ps20/gp116 and ps290/gp388 are homologs to *Acb1* and *Apyc1* , which could allow phage G to evade the host immune response by selectively degrading host cyclic nucleotide immune signals (Hobbs et al., 2022). Further functional investigation and validation of these antiviral escape genes are warranted.

Spore-related host regulatory proteins are expanded within phage G. The host of phage G is a *Lysinibacillus*, a gram-positive endospore former, with endospore formation being an ancient form of dormancy (Schwartz et al., 2023). Many phages have been identified to have spore-forming related homologs (Schwartz et al., 2023), but few, if any, have as many of the genes as phage G. Around five of the spore-related proteins were found amongst the viral particle proteomics. However, other spore-related proteins not found in particle proteomics may be expressed only during viral lytic infection inside the host. Phage G potentially evolved later than Moose phage W30-1, and appears to have acquired more spore related genes from its host *Lysinibacillus* sp. PGH. Encoding various prespore proteins within the virion particle may regulate the ability of the host to enter the spore stage at the point of infection. Multiple copies of the *RsfA*-like and *RsfA*-like without the full domain (ps281/gp379) within phage G may be repressors to stationary phase and/or spore formation (Wu & Errington, 2000). Spore protein D or small acid-soluble spore protein D with minor alpha/beta-type SASP is a *sspD*-like ORF encoded by ps168/gp266, that if like *sspD* like other SASPs would bind dsDNA and protect it from UV damage (Setlow et al., 2000). *YtxC*-like and stage V sporulation protein K are unknown transcriptional regulators of the sporulation; mutations in *spoVK/spoVJ* impair sporulation (Resnekov et al., 1995).

Methylation within phages provides higher-order regulation of DNA recognition, gene expression, replication, survival, and evasion of host response (Sun et al., 2020; Ding et al., 2023). Direct nanopore sequencing of phage G allows for *de novo* methylation detection without bisulfate chemical conversion due to the local change in current across the biological nanopore (Ni et al., 2019). The methylome is genome-wide within phage G but observable functionally, mainly within the baseplate-tail regulation region. Methylation positions 77050-78199 are within the baseplate protein where the long tail fibers attach. This upstream tail fiber attachment baseplate methylation may regulate the length of the tail fiber or attachment of tail fibers to the baseplate. Tail fiber regulation is critical to recognizing and interacting with the surface receptor of the host (Taslem Mourosi et al., 2022). *PncA*-like protein, potentially a pyrazinamidase/nicotinamidase/isochromatase hydrolase, is also highly methylated. The function of ps177/gp275 is unknown in phage G. However, *PncA* hydrolases (ps177/gp275) can function in a variety of ways, including drug resistance, or if it is a nicotinamidase, then that is used in recycling NAD+ (Shang et al., 2018; Pardee et al., 1971). The trimeric auto-transporter adhesins pectin lyase (DUF2807-like) function within phage G is unknown, but is highly conserved in multiple phyla of bacteria (Irfan et al., 2024). *YozC*-like ORF ps508/621 is a conserved hypothetical protein in *Bacillus* spp. nevertheless, its function is unknown ( O’Reilly et al., 2023). A computational and experimental protein-protein interaction assay suggested that *YozC* interacts with *AlaS* , an alanine-tRNA ligase gene, to adenylate cyclase *CyaA* , which suggests that methylation could regulate this process (O’Reilly et al., 2023).

Particle biophysical characterization of phage G included both temperature and pH stress. The UT_MUT had slightly less resilience and higher variance amongst replicates to temperature and pH than the ATCC/RAW_WT. Over >70 °C and high/low pH (pH = 3 and 11) resulted in zero plaques regardless of phage G variant. Our data suggests that phage G generally cannot handle significant rapid shifts in pH and temperature. This sensitivity may explain why the culture has been hard to maintain across multiple labs.

Phage G represents a marvel of phage biology that has remained in the shadows for over five decades. This data provides the blueprint for further investigations unlocking the genomic repertoire of phage G. Further study in a variety of avenues is needed to unlock more of the functions within phage G as well as other megaphages. Greater efforts are also needed to cultivate megaphages as model systems to unravel the phage world’s giants.

## Methods

### Sampling, DNA Extraction, Library Prep, and Sequencing

Phage G was amplified using dilute agarose gels as a modification of the traditional double agar overlay plaque assay method (Serwer et al., 2007; 2009). Phage particles were purified from double agar plaque assay with SM buffer and chloroform treatment; then, an aqueous layer was extracted from centrifugation at 5 min at 4,000 x g. Phage particles were precipitated with acetone from the aqueous layer (Soleimani-Delfan et al., 2021). Briefly, 1 part of phage particles (10^9^ PFU mL^-1^) was suspended in 4 parts acidified acetone (5.5 pH), shaken vigorously for 2 min, and centrifuged for 5 min at 4,000 x g. We precipitated 10 mL of total G phage stock. The supernatant was decanted, and the phage pellet was left to dry with phage pellets resuspended in SM buffer. The sample was DNA was extracted from all strains of G phage using the NEB (New England Biolabs, Ipswich, MA) Monarch HMW DNA Extraction Kit for Tissues (T3060), with a minor modification: RNase treatment was not performed. The libraries were prepared using the Oxford Nanopore Native Barcoding Kit 96 V14 (SQK-NBD114.96, Oxford, United Kingdom). DNA was sequenced using an Oxford Nanopore Promethion 2 Solo.

### Quality Control

Illumina 2500 data paired-end reads were filtered using Trimmomatic with sliding window parameters of 3 leading and trailing bases at a PHRED score of 20 and a minimum read length of 50 bases (Bolger et al., 2014). Adapters were detected and removed with ILLUMINACLIP using the TruSeq3 paired-end fasta, allowing two seed mismatches, palindrome alignment scores of 30, and an adapter clip threshold of 10 in accordance with the documentation. Raw Nanopore fast5 sequencing reads were processed with default filtering parameters from the provided MinKNOW sequencing software (v21.02.2) comprising a minimum quality score of 7, with real-time fast base-calling and fastq generation enabled. The resulting base called fastq files were used in subsequent downstream analyses.

### De novo genome assembly

*De novo* assembly was conducted on UT_WT and RIT_WT samples utilizing Unicycler (v.0.5.0) as they were exclusively Illumina HiSeq data (Wick et al., 2017). UT_MUT was also assembled in the same manner, as well as a hybrid Illumina/Nanopore *de novo* assembly approach using Trycycler (v0.5.4) as both data types were available (Wick et al., 2021). Briefly, the assessment of *k*-mer subassemblies from Unicycler’s implementation of SPAdes (v4.0) resulted in a best-selected k-mer size of 71, generating a single contig with no detectable dead ends in sequence (Bankevich et al., 2012). Contigs with low coverage (<10 reads) were removed, resulting in one anchor segment of 499,819 bases, subsequently used to form the final uniting sequence and did not circularize. CheckV (v1.01) was used to assess the quality and completeness of all G phage strains. CheckV accesses the Minimum Information about an Uncultivated Virus Genome (MIUViG) quality metrics (Roux et al., 2019). Genome statistics were evaluated using our in-house tool, RustyOmeStats (github.com/raw-lab/RustyOmeStats).

### Annotation, SNP, Methylome, and phylogenetic analysis

Annotation was completed using MetaCerberus (v1.4), Foldseek, HHpred, and SWORD with manual by-hand curation. Prodigal-gv was used within MetaCerberus to call ORFs (Camargo et al., 2024), for all downstream annotation processes. To avoid confusion with previous manuscripts, we have named the new ORFs from the corrected annotation ps1-668 based on ATCC_WT annotation, with the original gp (“gene product”) from the RWH draft genome to remind the same. For example, while the *terL* gene was rearranged to the beginning of the genome as gp1 or G_1 in GenBank in NCBI_WT, it is ps583/gp1 having both the corrected number based on the proper topology and arrangement of the genome (i.e., ps - ORF number) and the original gp number. We used SWORD to align NCBI_WT against ATCC_WT to obtain the original gp numbers in relation to the new ps ORF names. Origin of replication ( *OriC* ) and phage/host promoters were found using PhagePro and Ori-Finder 2022 (Sampaio et al., 2019; Dong et al., 2022), and default parameters were used. tRNAs and tmRNAs were annotated using tRNAscan-SE 2.0 and Aragorn (Laslett & Canback, 2004; Chan et al., 2021)

Methylome and single nucleotide polymorphisms (SNPs) were analyzed using DeepSignal2 (v0.1.3) and MUMmer4’s NUCmer (v4.0.0rc1). ATCC_WT provided the reference for all SNP analyses for the other variants. RIT_WT had significantly more SNPs than the other strains, MUMmer’s delta-filter script with –identity set to 8 and –length set to 1000 was used to filter the RIT_WT .delta output from NUCmer. Methylation was completed using DeepSignal2 using standard default measurements, and only high-quality methylations were used for downstream analysis.

ANI and phylogenetics analysis was completed using fastANI (v1.34), MAFFT (v7.520), and IQTree2 (v2.2.5) (Katoh & Standley, 2013; Jain et al., 2018; Minh et al., 2020). FastANI was run using the default parameters outlined within the readme. MAFFT alignments were completed using the local options -linsi with 1000 bootstraps for all trees using parameter -bb 1000. IQtree2 maximum likelihood trees were selected based on Modelfinder (Kalyaanamoorthy et al., 2017).

### Particle biophysical characteristics

A working stock of RAW_WT/ATCC_WT and UT_MUT of 10^9^ PFU mL^-1^ was used for all experiments. All temperature treatments were performed for 30 mins shaken at 250 RPM within a thermomixer. After temperature treatment, the samples were cooled on ice for 5 mins, then proceeded to plaque assay. For pH experiments, the desired pH was obtained and samples were exposed for 30 min. The pH was neutralized using either HCl or NaOH. Post pH exposure, the pH was neutralized to the control pH of 7 before undergoing a plaque assay. The experiment was repeated 4 times to determine the average PFUs for the results.

### Proteomic sample preparation

After estimating protein content with BCA assay, 20μg of proteins from G-ATCC, G-Mutant, and GLK samples, as well as half of the GSL, FGL, and GLS2 samples underwent in-solution digestion. Each sample was mixed with 55μL of UTT buffer (8 M urea and 10 mM DTT in 50 mM TEABC) in 1.5 mL microcentrifuge tube, incubated at 30°C with 500 rpm shaking for 60 min, followed by alkylation with 5μL of 250 mM iodoacetamide at 23°C for 60 min in the dark. After diluting urea to below 1M with 50 mM triethylammonium bicarbonate, 100μL of each sample was transferred to new microcentrifuge tube for proteolytic digestion by adding 0.4µg of Trypsin/LysC (Thermo Scientific, catalog# A40009) and incubating overnight at 37°C. Finally, digested peptides were acidified with 1% formic acid to terminate the proteolytic digestion.

### LC-MS/MS analysis

Digested peptides (500 ng) were loaded onto EvoTip trap columns (Evosep, EV2013), after column conditioning, equilibration and washing steps according to manufacturer instructions, with all centrifugations carried out for 60 s at 800 g. Peptides on the EvoTips were separated on 150μm x 15cm EASY-Spray column (PepMap^TM^ 2μm C18 beads, Thermo catalog # ES906) using Evosep One LC system (Evosep, Denmark). Peptides were eluted with a gradient up to 35% solvent B (0.1% formic acid in acetonitrile, solvent A: 0.1% formic acid in water) at flow rate of 0.5μL/min, using a 44-min gradient method and detected in positive ion mode with an Orbitrap Exploris 240 mass spectrometer (ThermoFisher). Data were acquired using either data-dependent acquisition (DDA, top 20) or data-independent acquisition (DIA, mass isolation window 24 m/z). Mass resolution was set at 60,000 resolution (at 200 m/z) for full MS scan and 30,000 for MS/MS scans. HCD collision energy was set at 30.

### Database search

The label-free raw data was processed and searched with Proteome Discoverer (PD, version 2.5.0.400, Thermo Fisher Scientific), using Sequest HT search engine applied and matched to the reference G phage genome, its host genome, and common MS contaminants (e.g., keratin, trypsin). Modifications to the searches included carbamidomethyl as a static modification on cysteines (+57.021 Da) and oxidation as a variable modification on methionines (+15.995 Da). In comparison, precursor mass tolerance was set as 10 ppm and fragment mass tolerance of 0.02 Da. Both data-dependent acquisition (DDA) and data-independent acquisition (DIA.) Mass spectrometry-based data was collected.The raw datafiles were analyzed using DIA-NN (version 1.8.1) with reference G-phage genome, its host genome, and common MS contaminants (e.g., keratin, trypsin). Settings included FASTA digest for library-free search, deep learning-based algorithms for spectra and retention times prediction, and critical parameters set at 15.0 for mass accuracy, 20.0 for MS1 accuracy, and 4 for scan window. Enzyme specificity was trypsin with allowance for one missed cleavage. Carbamidomethyl modification on cysteine residues was fixed, and a match between runs (MBR) was enabled. Protein inference grouped on genes using a neural network classifier, quantification optimized for LC accuracy, cross-run normalization tailored to retention time-dependent dynamics, and smart profiling techniques for library profiling. Default settings were used for other parameters for comprehensive analysis.

## Data and Code Availability

The code used in this work are publicly available on GitHub (https://github.com/raw-lab/) All phage genomes used in this study are available on NCBI GenBank. Phage proteomic data is available on PRIDE.

## Funding

Andra Buchan, Stephanie Wiedman, Kevin Lambirth, Madeline Bellanger-Perry, Jose L. Figueroa III, and R.A. White III are supported by the UNC Charlotte Department Bioinformatics and Genomics start-up package from the North Carolina Research Campus in Kannapolis, NC.

## Acknowledgments

We also acknowledge the University Research Computing and the College of Computing and Informatics for computational and logistical support. We must further acknowledge Steven C. Hardies for his help with annotation of phage G and highly useful discussions.

## Conflicts of Interest

The authors declare no conflicts of interest. RAW is the CEO of RAW Molecular Systems (RAW), LLC, but no financial, IP, or others from RAW LLC were used or contributed to the study.

## Supplemental materials

**Fig S1:**Sequence logo plots for the long terminal repeat (LTRs) sequences comparing all phage G variants and Moose W30-1. a) All phage G variants b) All phage G variants with Moose W30-1.

**Fig S2:**Maximum likelihood trees for dihydrofolate reductase protein (ps30/gp125), Stage V sporulation protein K (ps483/gp596), *PhoH-like* protein (ps65/gp162). Alignments were completed with MAFFT then maximum likelihood trees with 1000 bootstraps. T4 and T5 phages were used as an outgroup. AlphaFold2 protein predictions are showcased for various representative clades including phage G.

**Fig S3**. *OriC* plot detection from Ori-Finder 2022. Accessed Aug 4th 2024 (https://tubic.org/Ori-Finder/)

**Table S1**.This includes many tables of various supplemental data as tabs. Tab 1 is the ICTV official phage taxonomy from TaxMyPHAGE. Tab 2 is the hand curated annotation for phage G using ATCC_WT as the main reference. Tab 3 is the hand curated annotation for Moose phage W30-1. Tab 4 is the SNP table for phage G variants across all using MUMmer 4.0. Tab 5 is all the predicted promoters from phage G (ATCC_WT) both host/phage from PhagePro. Tab 6 is the tRNAs and tmRNA predictions from tRNAscan-SE and Aragon.

## Notes

### Competing Interest Statement

The authors have declared no competing interest.

