## Supplementary figures and images for "Unlocking the genomic repertoire of a cultivated megaphage"

### Fig S1

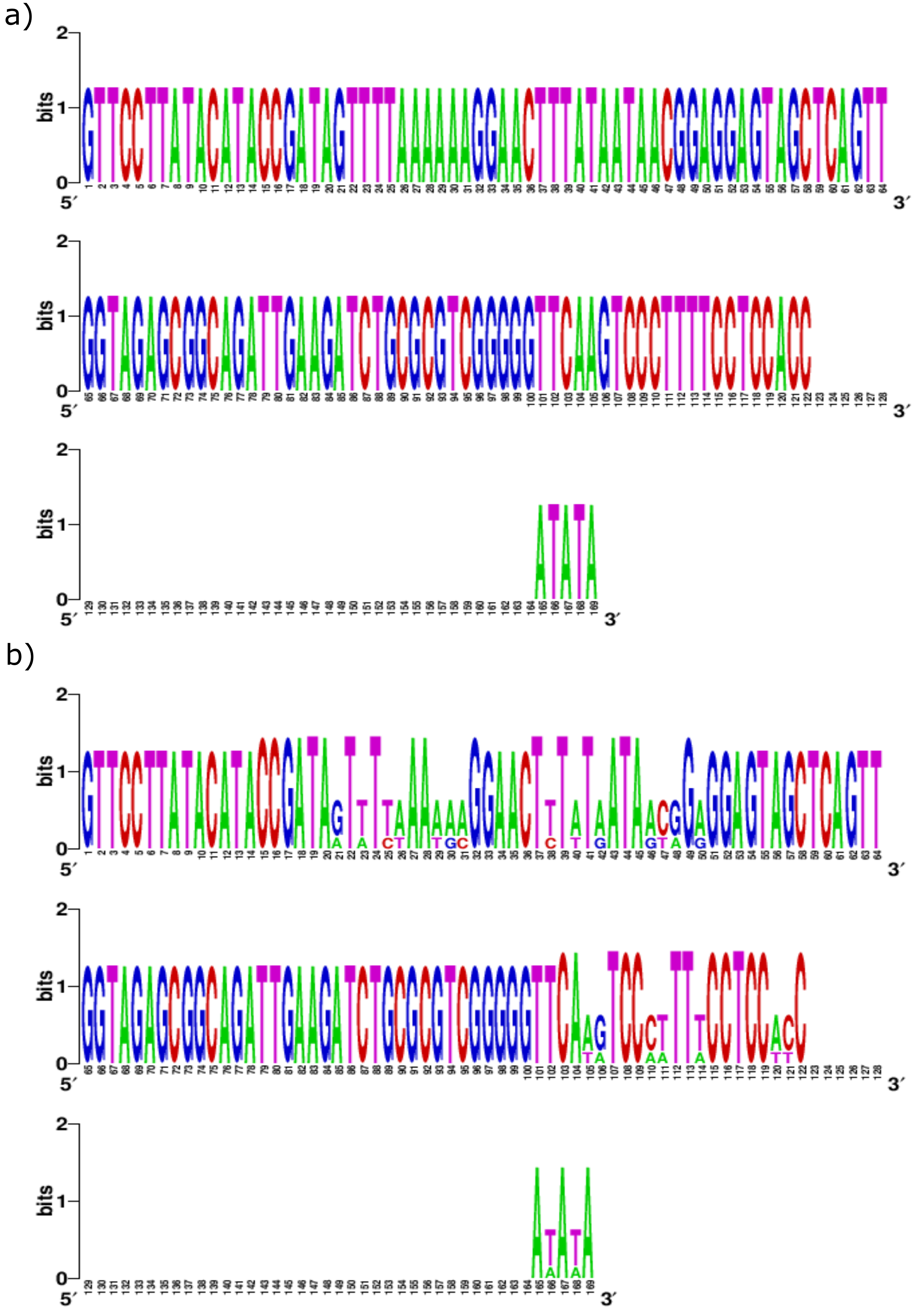

### Fig S2

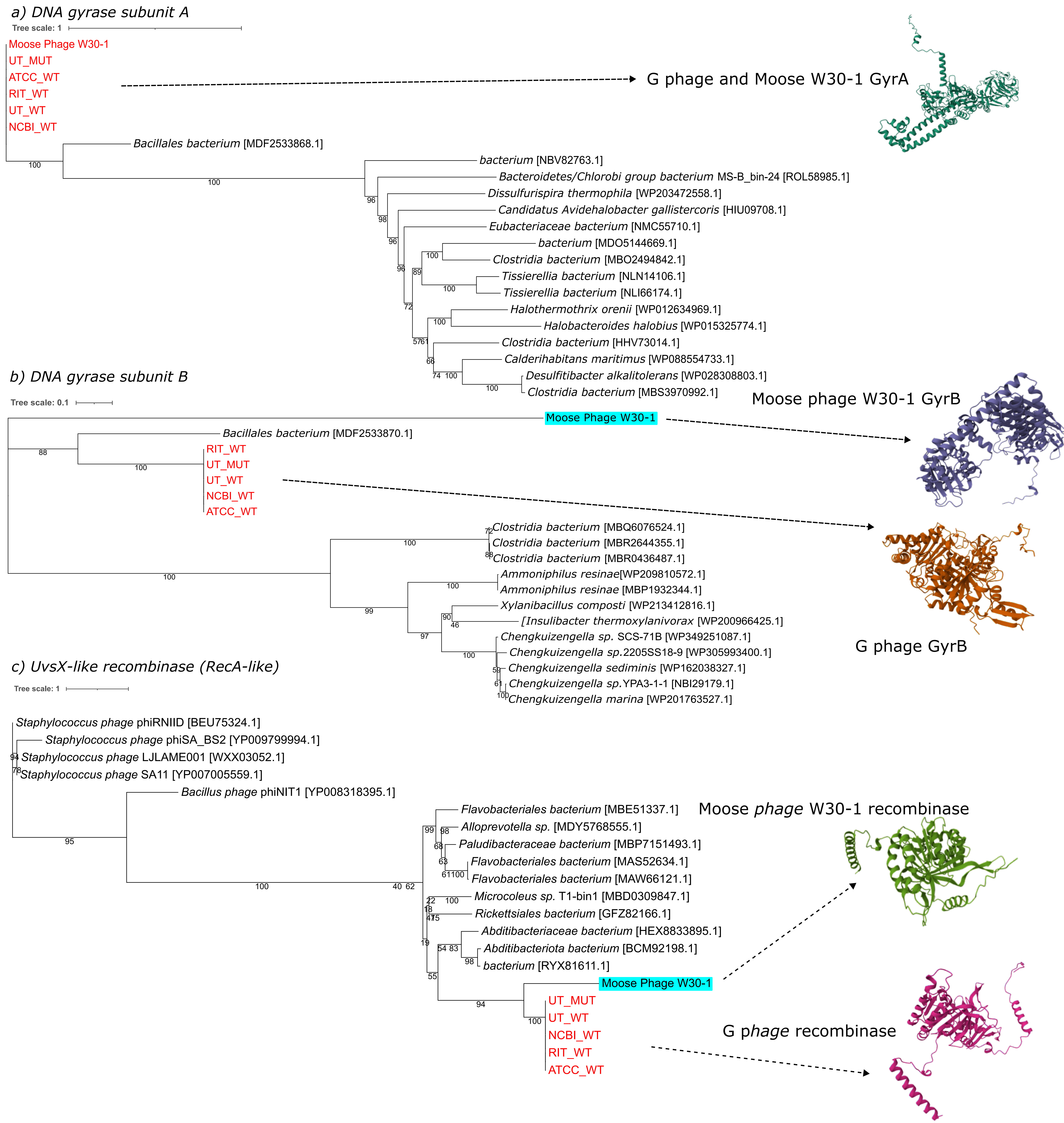

### Fig S4

#### a) Dihydrofolate Reductase Protein

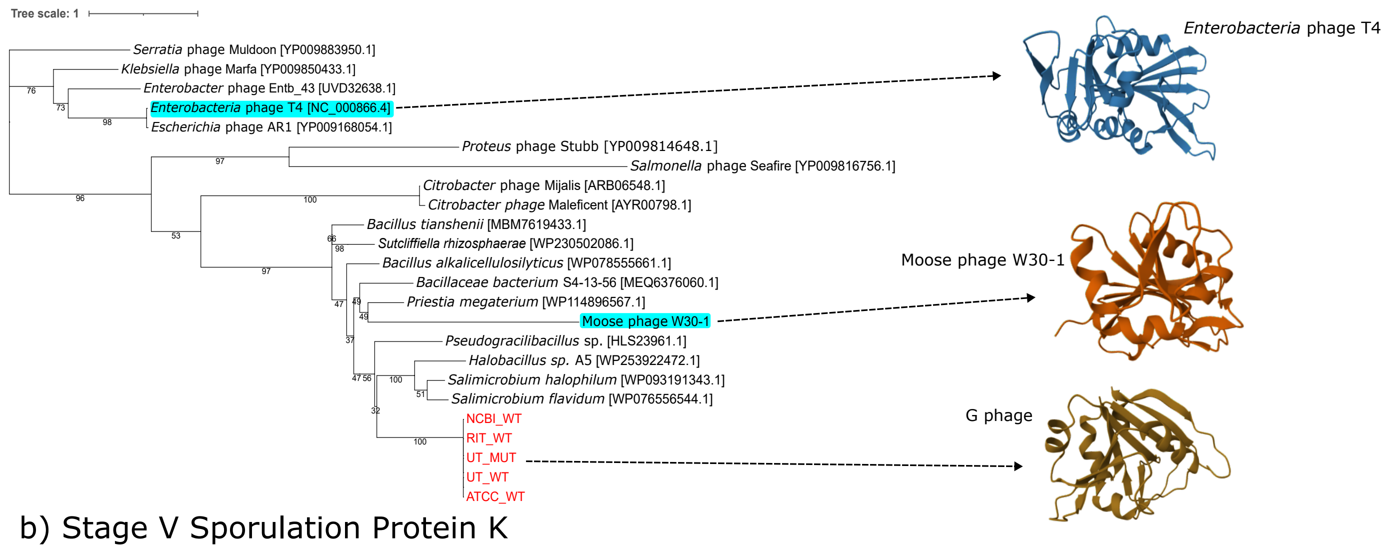


#### b) Stage V Sporulation Protein K

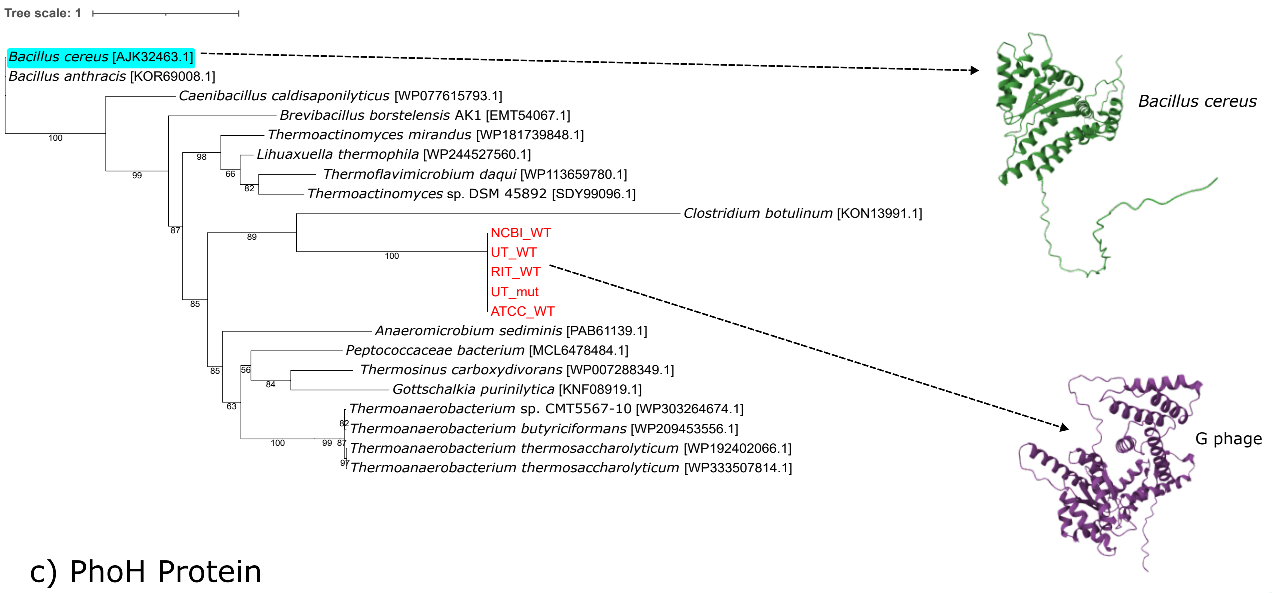


#### c) PhoH Protein

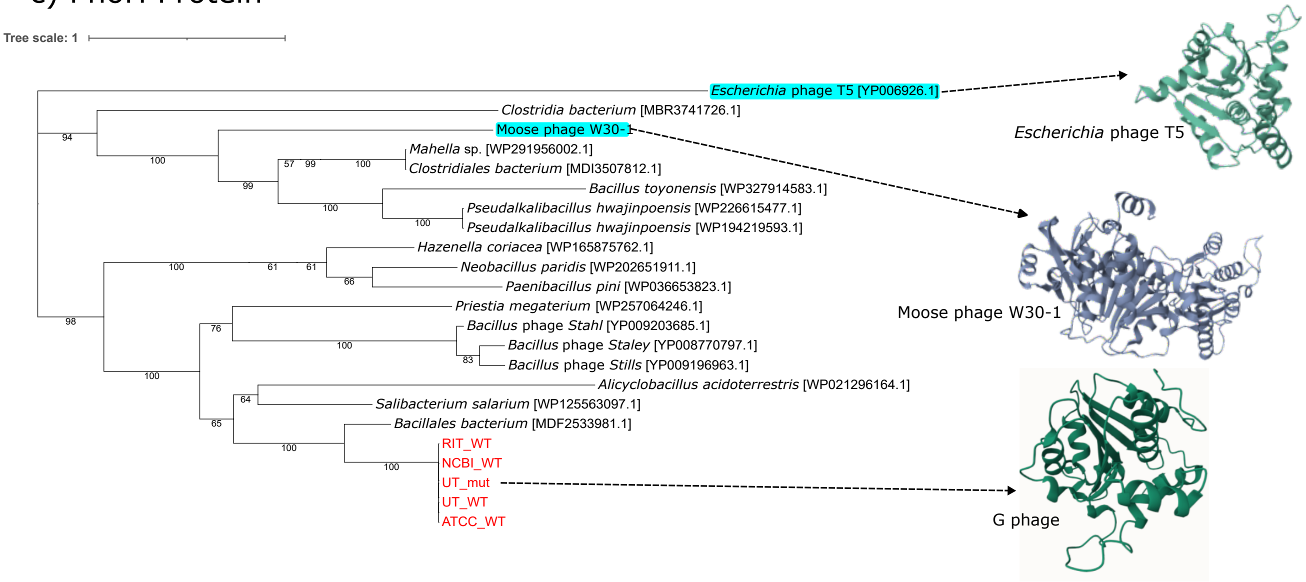

### Fig S5

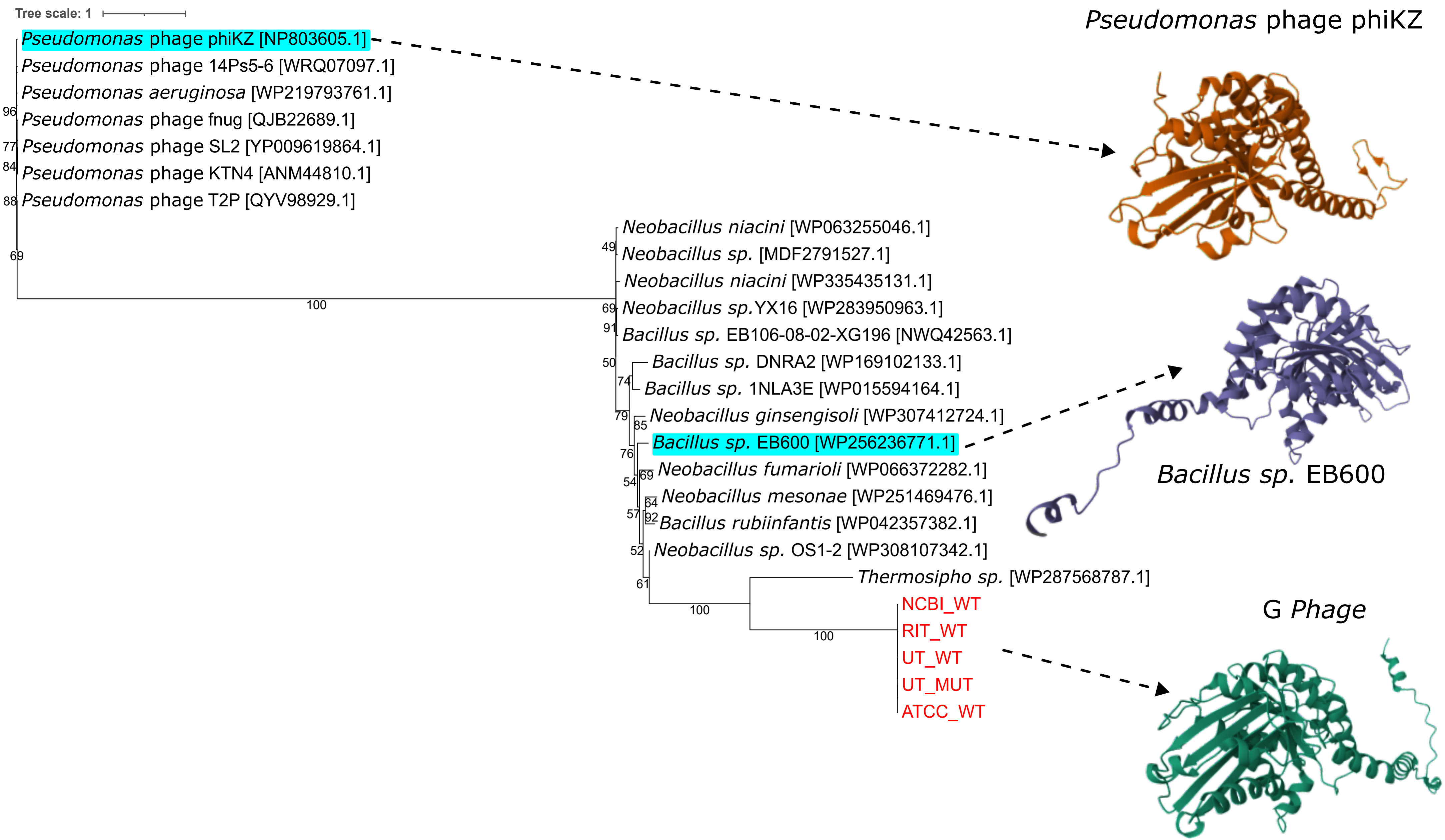
