## Supplementary material for "Unlocking the genomic repertoire of a cultivated megaphage": Fig S3

| RefSeq | ATCC_WT |
| --- | --- |
| OriC length | 406 nt |
| OriC AT content | 0.52 |
| The location of oriC region | 1..406 nt |
| The extremes of GC disparity | 15 nt (minimum), 499918 nt (maximum) |

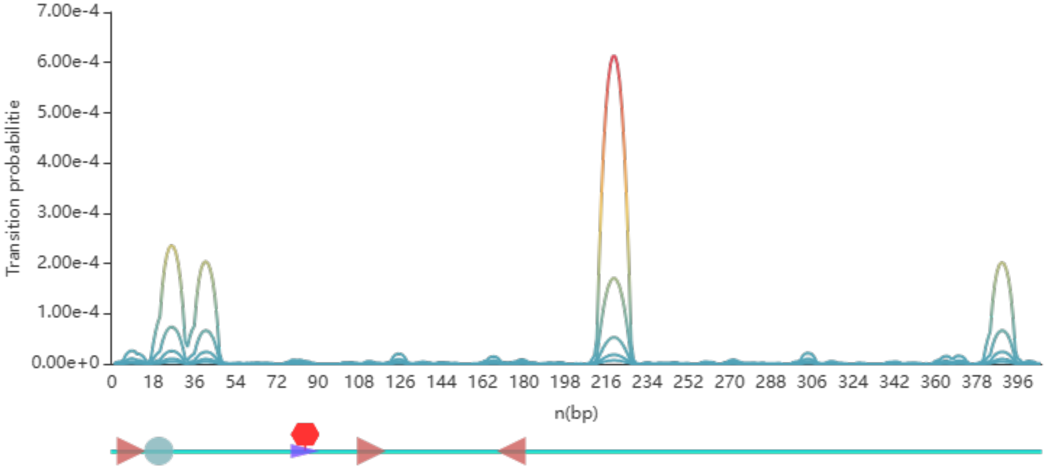


DnaA box · DnaA-trio · ATP-DnaA box · GATC · CtrA binding motif · Fis binding motif · IHF binding motif

-GTTCCTTATACATACCGATAGTTTTAAAAAAGGAACTTTATAATAACGGAGGAGTAGCTCAGTTGGTAGAGCGGCA
GATTGAAGATCTGCGCGTCGGGGGTTCAAGTCCCTTTTCCTCCACCATATATGGGGAGATAGACCGTAAGTGGAAGC
GGGCAGGACTGTAAATCCTGTGAGAAATCCTTGGGTGTTCGATTCGCCCTCTCCCCACCATTTTATTAATTATGGCG
CGTTAGCCAAGTGGTAAGGCATCCGGCTTTCTACCGGAGATTACACGAGTTCGAACCTCGTACGCGCTACCATTTTT
GCCGGTTTAGCTCAGCGGTAGAGCGCTCCCCTTGTAAGGGAGATGTCGTGGGTTCAAATCCTATAACCGGCTCCATG
TTATTTTAAAGGAGGAATTCGC-

5'-DnaA box-3' · 3'-DnaA box-5' · Spacer · DnaA-trio · ATP-DnaA box
AT-rich region · GATC · CtrA binding motif · Fis binding site · IHF binding site

| RefSeq | ATCC_WT |
| --- | --- |
| OriC length | 376 nt |
| OriC AT content | 0.58 |
| The location of oriC region | 611..986 nt |
| The extremes of GC disparity | 15 nt (minimum), 499918 nt (maximum) |

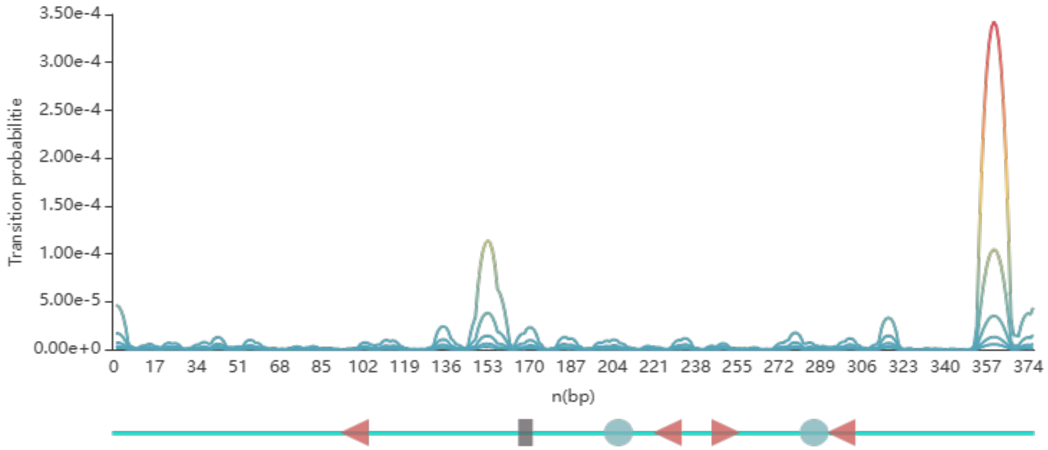


DnaA box · DnaA-trio · ATP-DnaA box · GATC · CtrA binding motif · Fis binding motif · IHF binding motif

-TATTTGCAGGTTGTAAGTGGAATGATAGCCACTAGAATCCATATAGCACAACGAAAACATGAGGTAGCTCCTCATT
CGTCTTAGGTGAACCTCCCTGTGCAAAAGAGAGTTTGATTCTCTCCACCTGCTCCATATATGTCATCGGTAGTTTAA
ATAGAATAGGCTTAGATAATCTGCGACCAAAAGATAGTGGACCTAAGTTCTAAGAACAGTCCCATTGTGATGGGGAA
AGTTAAGGTGCGATTCCTTATCGAGACACCATTCGGAGGGTTACTCTAATTGGTAAGAGAACAGTCTTGAAAACTGT
CGTAGGTATAAAAGCCGATGGGGGTTCGAGTCCCTCACCCTCCGCCATATAATTTATTAGGAGAAATCT-

5'-DnaA box-3' · 3'-DnaA box-5' · Spacer · DnaA-trio · ATP-DnaA box
AT-rich region · GATC · CtrA binding motif · Fis binding site · IHF binding site
